# Micro-topographical guidance of macropinocytic signaling patches

**DOI:** 10.1101/2020.08.29.273185

**Authors:** Gen Honda, Nen Saito, Taihei Fujimori, Hidenori Hashimura, Mitsuru J. Nakamura, Akihiko Nakajima, Satoshi Sawai

## Abstract

In fast moving cells such as amoeba and immune cells, spatio-temporal regulation of dendritic actin filaments shapes large-scale plasma membrane protrusions. Despite the importance in migration as well as in particle and liquid ingestion, how these processes are affected by the micrometer-scale surface features is poorly understood. Here, through quantitative imaging analysis of *Dictyostelium* on micro-fabricated surfaces, we show that there is a distinct mode of topographically guided cell migration ‘phagotaxis’ directed by the macropinocytic Ras/PI3K signaling patches. The topography guidance was PI3K-dependent and involved nucleation of a patch at the convex curved surface and confinement at the concave surface. Due to the topography-dependence, constitutive cup formation for liquid uptake in the axenic strain is also destined to trace large surface features. Given the fact that PI3K-dependency of phagocytosis are restricted to large particles in both *Dictyostelium* and immune cells, topography-dependency and the dual-use of membrane cups may be wide-spread.

## Introduction

Large-scale deformation of plasma membrane during cell migration, particle and liquid ingestion depends on physical cues such as substrate rigidity and topography (Champion and Mitragotri, 2006; Clarke et al., 2010; Kim et al., 2009; Rajnicek et al., 1997; Ray et al., 2017; Teixeira et al., 2006). Cell migration along surface structures such as ridges and grooves, is generally referred to as contact guidance and thought to play pivotal roles in neural development (Reig et al., 2014), tissue repair, immune response and cancer invasion (Friedl and Alexander, 2011). Nano- and micro-fabricated platforms have clarified how geometrical constraint affects focal adhesions distribution and actin stress fibers (Mathur et al., 2012; Oakley and Brunette, 1993; Ray et al., 2017). Focal complexes, substrate-anchoring clusters containing ECM-bound integrins that engage vinculin and talin associated with actin stress fibers, are restricted to be distributed within ridges and grooves (Franco et al., 2011; Ray et al., 2017). There alignment of stress fibers plays a major role in generating anisotropic contractility. While such a mechanism appears to be wide-spread in cells of epithelial or mesenchymal nature (Franco et al., 2011; Oakley and Brunette, 1993; Ray et al., 2017), topographical guidance in fast-moving amoeboid cells (Driscoll et al., 2014; Kwon et al., 2012; Sun et al., 2015; Wilkinson et al., 1982) which do not have stress-fiber and can migrate independently of cell-substrate adhesion (Lämmermann et al., 2008) is far less understood. Neutrophils are known to elongate along a few micron square grooves of a hemocytometer surface (Wilkinson et al., 1982). T-cells migrate along parallel ridges/grooves whose widths are hundreds of nanometers (Kwon et al., 2012). Adhesion-independent mode of migration in T-cells occurs under 2D confinement only if there is topographical asymmetry in the physical surrounding (Reversat et al., 2020). Macrophages are also known to spread along ridges and grooves (Wójciak-Stothard et al., 1996). Besides contractility, the other dominant determinant of directionality in fast-migrating amoeboid cells is the leading edge protrusion. While there are large body of work addressing how diffusible chemoattractants determine when and where the leading edge forms to steer the cells, how they are guided by topography remains largely unknown (Sales et al., 2017).

The leading protrusion formed during cell migration has a large overlap in its molecular compositions with those formed during particle and liquid ingestion and thus the distinction between these processes are sometimes obscure (Heinrich and Lee, 2011). Conventionally, uptake of particle and liquid are referred to as phagocytosis and macropinocytosis, respectively. As in the leading edge of migrating cells, macropinocytosis and phagocytosis involve large-scale conversion from contractile actomyosin to protrusive branched actin meshworks that requires activation of the Arp2/3 complex for the side-branching nucleation (Molinie and Gautreau, 2018; Pollard, 2007). The resulting actin polymerization generates the protruding force for the expanding edge of a cup-shaped membrane invagination for ingestion (Jaumouillé et al., 2019; Rougerie et al., 2013). Phagocytosis often refers to specific adhesion-dependent engulfment in immune cells, where the surface of the ingesting particle is decorated with opsonins; i.e. scaffold antigens or complements which through membrane-bound receptor signaling (Swanson, 2008) processively extends the protruding edge of the cup along the attached solid surface (Case and Waterman, 2015; Jaumouillé and Waterman, 2020). Macropinocytosis on the other hand refers to a self-organizing process where the shaping of the membrane by the branched actin meshworks does not require a solid surface (Swanson, 2008) and can occur constitutively (Williams and Kay, 2018). While this property makes it suitable for the uptake of nutrient media as well-known in cancer cells and *Dictyostelium*, macropinocytic particle uptake is also known for the entry of pathogenic bacteria into the host cells (Amara and Mercer, 2015). Non-opsonized polystyrene beads can also be ingested by amoeba *Dictyostelium* as well as macrophages and dendritic cells (Gilberti and Knecht, 2015; Mu et al., 2018; Pacheco et al., 2013). The receptor-independent cues that guide these macropinocytic/phagocytic membrane protrusion are still poorly understood.

Unlike endocytic cups mediated by clathrin, caveolin and BAR domain-containing proteins whose nanometer-scale topography dependence have been well studied (Galic et al., 2012; Zhao et al., 2017), phagocytic and macropinocytic cups involve global reorganization of actin cytoskeletons. In *Dictyostelium*, early organization of phagocytic/macropinocytic cup formation begins with the appearance of micrometer-size patches enriched in dendritic actin filaments whose inner-domain is characterized by strong accumulation of phosphatidylinositol (3,4,5)-trisphosphate (PIP3) (Hoeller et al., 2013) along with GTP-bound form of Ras and Rac. Depolymerizing factor coronin is distributed further into the cytoplasmic side (Gerisch, 2010; Hacker et al., 1997; Maniak et al., 1995). These active signaling patches are self-amplified by a positive feedback loop involving Ras and PI3K (Fukushima et al., 2019; Sasaki et al., 2007; Taniguchi et al., 2013) and serve as a common precursor or ‘template’ for phagocytic/macropinocytic cup formation (Gerisch et al., 2009; Veltman et al., 2016). As the patch increase in size, its outer edge enriched in the SCAR/WAVE complex protrudes outward to form a circular ruffle (Veltman et al., 2016). Loss of RasGAP NF1 or IqgC enhances both phagocytosis and macropinocytosis (Bloomfield et al., 2015; Marinović et al., 2019; Williams and Kay, 2018) indicating that Ras act positively on both processes. While deletion or pharmacological inhibition of PI3K suppresses liquid uptake, deleterious effect on phagocytosis is limited to the uptake of large particles (Buczynski et al., 1997; Chen et al., 2012; Hoeller et al., 2013). Patches with identical molecular organization are observed in the ventral side facing the substrate where they appear as traveling waves (Asano et al., 2008; Bretschneider et al., 2009; Brzeska et al., 2016, 2014; Taniguchi et al., 2013; Veltman et al., 2016). The ventral patches are thus thought to be a frustrated form of a macropinocytic/phagocytic cup (Gerisch, 2010; Gerisch et al., 2009) similar to the frustrated phagocytosis in macrophage placed on an opsonized surface (Barger et al., 2019; Masters et al., 2016). In this work, to clarify the relationship between the surface microscale topography and actin patch initiation, propagation and termination, live-cell imaging analysis of *Dictyostelium* on micro-fabricated surfaces was performed. We demonstrate that the propagating ventral patches guide cell migration along a micrometer-scale ridge in a PI3K-dependent manner. Quantitative analysis shows that the nucleation of the patches occurs at the convex surface whereas its propagation is restricted at the concave surface. Our results suggest that these properties allow the macropinocytic cup to engulf extracellular fluid by default while at the same time directing it to faithfully trace bent and bifurcating ridges when in contact with structured surfaces.

## RESULTS

### Ventral actin patches propagate along microridges and orient polarized AX4 cells

In order to first gain an over-view of the micro-topography dependency of *Dictyostelium* cell deformation, we studied an aggregation-stage axenic strain (AX4) expressing GFP-Lifeact on an SU-8 structured substrate with straight ridges. Here, the ridges employed were 1 μm in height, 3 μm in width and placed in parallel at intervals of 3 μm (see Materials and Methods). Based on time-lapse confocal imaging, cells were manually scored at each time frame for the presence of intense patches of F-actin on the ventral plasma membrane. On both flat and structured surfaces, the percentage of the cells that exhibited the patch increased from two hours after plating (Fig. S1A). The maximum percentage of patch-positive cells were 79 ± 13 % and 98 ± 1 % for flat and micro-structured surfaces, respectively (Fig. S1A, right panel, 250 min). Cellular movement on a flat surface was almost isotropic in direction (Fig. 1A, Fig. S1B). In the presence of ventral F-actin patches, cells showed relatively small net displacement compared to those without (Fig. S1B). Actin patches propagated along the ventral plasma membrane and the area in contact with the substrate expanded as the front traveled outward at the edge (Fig. 1B, Movie. S1). The direction of patch propagation frequently changed as shown in Fig. 1B (red trajectories). On the other hand, cells on the micro-structured surface migrated persistently along the ridges in the presence of patches (Fig. 1C, Fig. S1C). As in patches found on flat surfaces (Asano et al., 2008; Bretschneider et al., 2009), F-actin was most densely accumulated at the outermost edge of the patch which surrounds the inner territory enriched in phosphatidylinositol(3,4,5)-trisphosphate (PIP_3_) (Fig.1D, see also Supplementary Info Fig. S3A-B). The cells were locked-in to a ridge and their movement seldom deviated from a single linear track. The actin patches remained in the cell anterior as the cell moved forward along the ridge (Fig. 1D, Movie. S2). More than 80% of migratory direction were oriented parallel to the ridges in patch-positive cells on the micro-structured surface (Fig 1E). In contrast, the speed of cells was not largely affected (Fig. 1F). On a flat surface, the persistence time of cell migration was 0.54 min (N = 23 cells) in the presence of ventral patches, compared to 3.0 min (N = 36 cells) in the absence of patches (Fig. 1G; Flat, patch(+), Flat, patch(-)). On the structured surface, persistence time 13.7 min of migratory direction in the presence of ventral patches (N = 34 cells) was three times higher compared to 4.6 min in the absence (N = 21 cells) (Fig. 1G; Struc., patch(+), Struc., patch(-)).

**Figure 1.**
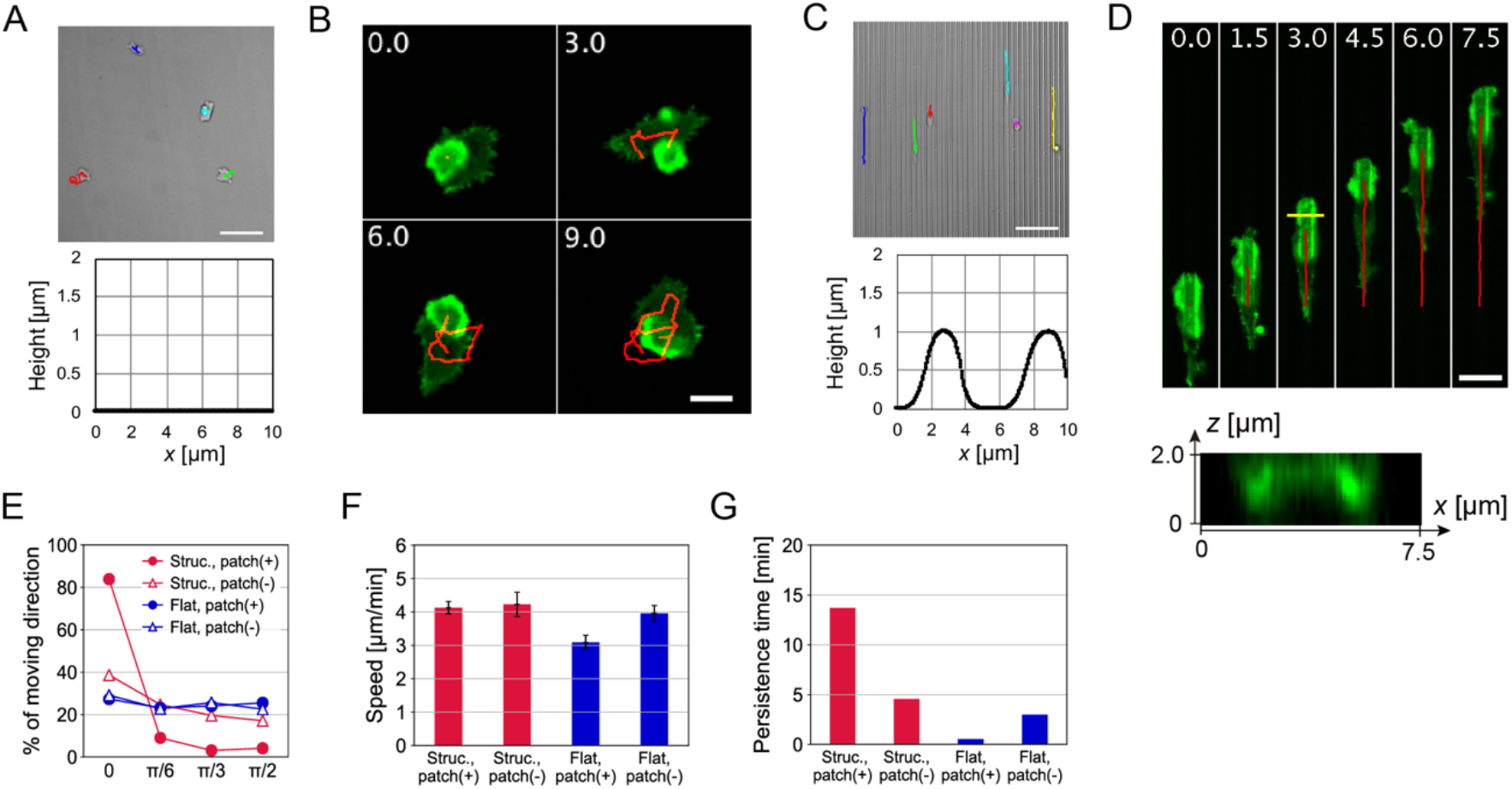
Guidance of ventral actin patches and membrane evagination in *Dictyostelium* AX4 cells on a microridged surface. (**A-D**) Cell trajectories and the ventral F-actin patch dynamics in aggregation-stage AX4 cells on non-structured (**A** and **B**) and microstructured SU-8 surfaces (**C** and **D**). (**A** and **C**, upper panels) Transmitted light images of a representative field of view. Colored lines: trajectories of individual cells for 20 min. Scale bars, 50 μm. (**A** and **C**, lower panels) Representative surface geometry. (**B** and **D**) Time-lapse confocal images of the F-actin patch. Green: GFP-Lifeact fluorescence; *z*-slice near the SU-8 surface (*z* = 0) (**B**) and maximum intensity projection (MIP) from *z* = 0 to 2 μm (**D**, upper panel) and the cross-section along the yellow line (**D**, lower panel). Red lines: centroid trajectories of the F-actin patch. Time in minutes. Scale bars, 10 μm. (**E**) Angular distribution of cell centroid displacement relative to the ridge direction. (±): the presence or absence of the ventral F-actin patch. (**F**) Cell migration speeds (mean ± s.e., N = 34, 21, 23, 36 cells). (**G**) Persistence time of cell displacement.

The F-actin patch and the leading edge traveled along a single ridge and rarely traversed to neighboring ridges. While some cells migrated in one direction for over 30 minutes before switching to the opposite direction, others frequently made turns and consequently showed small net displacement; less than 80 μm for 50 min (Fig. S1D). Change in the direction of cell migration was accompanied by either patch reversal (Fig. S1E) or splitting (Fig. S1F). In patch reversal, the actin patch starting from the cell anterior traveled to the opposite end (Fig. S1E). In patch splitting, the anterior actin patch split in half and a daughter patch reached the posterior end and became a new front while the other patch disappeared (Fig. S1F). Cells that exhibited frequent patch reversal and splitting showed small net displacement (Fig. S1G). While these observations indicate strong correlation between direction of cell movement and patch propagation, persistent migration was rarely observed in growth-stage cells. There, almost the entire ventral side of the plasma membrane was covered by a single continuous patch or a few separate patches, resulting in large ruffles projected in many directions (Fig. S2A, Flat). Small patches were restricted at the ridges (Fig. S2A, Ridge, 24-48 sec) while larger ones often covered several ridges (Fig. S2A, Ridge, 72-120 sec). Regardless of the presence of actin patches, persistent migration along the microridge rarely occurred (Fig. S2B,C, compare to Fig. 1E) indicating that micro-topograhical features guide patch propagation but requires additional cell polarity for persistent migration.

These observations raise a question whether the F-actin/PIP3 patch formation which is thought to be a constitutive process that serves as a precursor for the macropinocytic/phagocytic cup formation (Gerisch et al., 2009; Veltman et al., 2016) was replaced by another distinct process when presented with curved surfaces. A previous work has shown that, in growth-stage or cells early into differentiation, the F-actin patches are extinguished by treating the cells with PI3kinase inhibitor LY294,002 (Taniguchi et al., 2013). We found that in aggregation-stage cells too, F-actin patches are extinguished with LY294,002 treatment in a dose-dependent manner (Fig. 2A). When cells migrating along the SU-8 ridge is applied locally with LY294,002 using a microneedle, F-actin patches disappeared immediately and the cell trajectories began to deviate from the ridge (Fig. 2B-C). Directional bias decreased to a level comparable to non-treated cells without the ventral patch (Fig. 2D, Fig. 1E; Struc., patch(-)). A mock treatment neither extinguished patches nor impaired the topographic guidance (Fig. 2D). The results indicate that the micro-topographic guidance is PI3kinase-dependent and thus distinct from PI3kinase-independent, biased cell migration along much finer submicrometer-scale ridges (Sun et al., 2015). Spatial organization of the molecular components was also indistinguishable (Fig. S3) from those known for the ventral patches on flat surfaces (Schroth-Diez et al., 2009; Taniguchi et al., 2013; Veltman et al., 2016).

**Figure 2.**
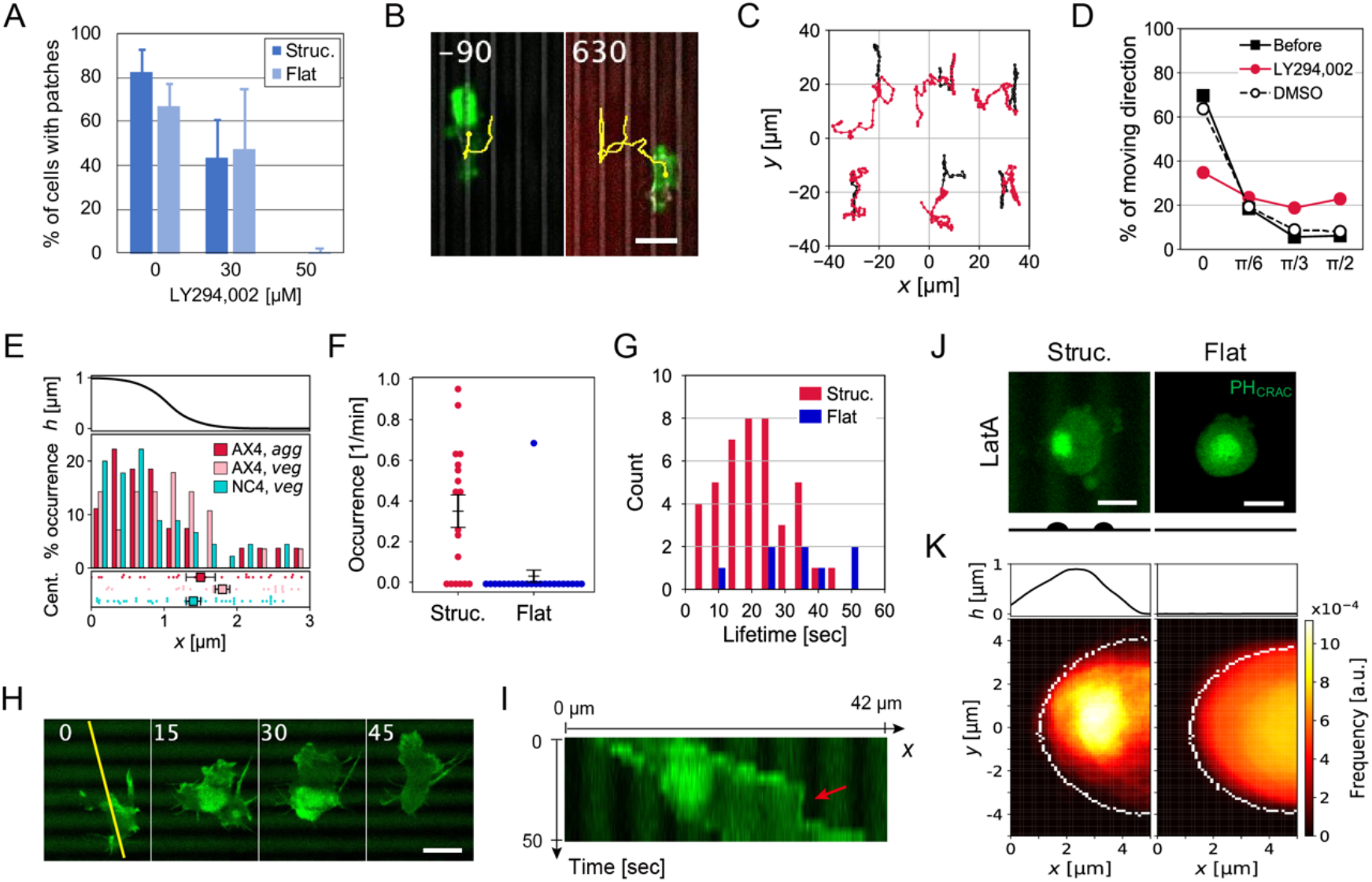
Micro-topographical guidance is PI3K-dependent. (**A**) Fraction of patch-positive AX4 cells in aggregation-stage after LY294,002 treatment (mean ± s.e., 13 cells per condition). Averaged for 15 min, and compared in the same field of view before (*t* = −15 to 0 min) and after (*t* = 10 to 25 min) application of the inhibitor (*t* = 0 min). The measurement was from several field of views in two independent trials per condition. (**B**) Merged confocal images (green: GFP-Lifeact fluorescence, red: Alexa594 in the LY294,002 solution, grey: transmitted light, yellow lines: the trajectory of cell centroid). Time in sec; LY294,002 application from a microneedle at t = 0. Scale bar, 10 μm. (**C**) Cell trajectories from t = −5 to 0 min (black) and from t = 0 to 13 min (red). (**D**) Angular distribution of cell displacement relative to the ridges, before (black solid line; N = 14 cells) and after LY294,002 application (red line; N = 6 cells). DMSO mock control (black broken line; N = 8 cells). (**E**) Distribution of patch nucleation along the x-axis (upper panel: the ridge z-profile) for aggregation-stage (*agg*) AX4 cells, vegetative (*veg*) AX4 and NC4 cells (middle panel). Cell position at the time of patch nucleation (bottom panel, mean ± s.e., N = 27 (AX4, *agg*), 28 (AX4, *veg*) and 45 patches (NC4, *veg*), each dot represents a unique cell). (**F**) Frequency of ventral F-actin patch nucleation in vegetative NC4 cells (mean ± s.e., N = 18 (Structured) and 23 cells (Flat), each dot presents a unique cell). (**G**) Lifetime distribution of the F-actin patches in vegetative NC4 cells. (H) Lifeact-GFP/NC4 on microridges (green: Lifeact-GFP fluorescence; MIP from *z* = 0 to 3 μm). Time in sec. Scale bar, 10 μm. (**I**) A kymograph along the yellow line in (H). The image is enlarged eight times in time-axis. (**J**) Representative snapshots from confocal images of vegetative NC4 cells expressing PH_CRAC_-GFP that are treated with 3 μM LatA on microstructured (left) and non-structured (right) surfaces (green: PH_CRAC_-GFP fluorescence; MIP from *z* = 0 to 3 μm, the lower schematic indicates ridge positions). Scale bars, 5 μm. (**K**) The average spatial profile of ventral PH_CRAC_-GFP fluorescence in LatA-treated cells; microstructured (left, N= 15 cells) and non-structured (right, N = 14 cells) surfaces. The ridge profile is shown for reference in the upper panel. White borders indicate the average cell contours. See Materials and Methods for details.

### PI3K signaling is induced by microtopography

In order to see whether the micro-topography potentiates the appearance and lifetime of the patches, we first counted the positions of patch nucleation relative to the surface topography. The result shows that most ventral actin patches were initiated at the ridge in both growth- and aggregation-stage cells (Fig. 2E; “AX4, *veg*”, N = 28 patches; “AX4, *agg*”, N = 27 patches), independent of the position of the cell centroids (Fig. 2E, bottom). The duration of the patch-positive phase in growth-stage cells was on average 244 ± 54 sec (N = 20 events) and 654 ± 111 sec (N = 24 events) on flat and ridged surfaces, respectively. In aggregation-stage cells, majority of patches persisted throughout our timelapse observations (50 min). Conversely, when the surface was coated with lectin wheat germ agglutinin (WGA), which promotes attachment of *Dictyostelium* cells to the substrates (Yoshida et al., 1984), the occurrence of actin patches decreased markedly (Fig. S4A, B) and lifetime decreased to 4.0 ± 0.6 min (structured, N = 22 patches; flat: no patch data), and the directional bias also diminished (Fig. S4C, compare to Fig. 1E). An earlier study has shown that the signaling patches that appear in the ventral membrane are the results of exaggerated Ras activity in the axenic cell-lines due to its null-mutation in RasGAP NF1 (Veltman et al., 2016). Therefore we tested the patch properties in the parental non-axenic NC4 strain which has the intact RasGAP. On a flat non-structured surface, only a few percent of NC4 cells exhibited ventral F-actin patch (4.3 %, N = 23 cells), in agreement with the recent report (Veltman et al., 2016). In contrast, we found that the percentage increased drastically on a microridged surface (67 %, N = 18 cells). The occurrence of patches were 0.35 ± 0.08 /min and 0.03 ± 0.03 /min for structured and non-structured surfaces, respectively (Fig. 2F). Moreover, majority of the patches were initiated at the ridge (Fig. 2E, “NC4”, N = 45 patches), indicating that the surface ridge geometry serves as an essential trigger for the patch initiation in NC4 cells. These patches were relatively short-lived (Fig. 2G, Struc.: 21 ± 2 sec, N = 42 patches, Flat: 34 ± 5 sec, N = 8 patches) and did not facilitate migration, however competed with the leading edge extension (Fig. 2H; t = 15 ~ 30 sec, Fig. 2I; red arrow). In aggregation-stage NC4 cells, F-actin patches were not detected under our conditions. These results indicate that both the onset and the lifetime of F-actin patches are topography-dependent.

To further study the nature of topographical dependency, we employed NC4 expressing PH_CRAC_-GFP as a marker for PIP3. Surprisingly, in the LatA-treated growth-stage NC4 cells, the patch in the ventral plasma membrane was already present on the flat surface (Fig. 2J, right, Fig. 2K, right; N = 14 cells) in addition to the structured surface (Fig. 2J, left). This stands in contrast to the absence of patches on the flat surface in the untreated cells (Fig. 2F). On the structured surface, LatA-treated cells were dislodged from the ridge and usually found between the two neighboring ridges, and the PIP3 patches at the ventral-side were initiated above the ridge (Fig. 2J, left) which is consistent with the ridge-dependence in non-treated cells (Fig. 2F). These patches remained at the site of initiation and disappeared spontaneously (Movie. S3). The frequency map of the PH_CRAC_-GFP patches along the ventral-side of the plasma membrane shows a strong bias in their occurrence at the ridge (Fig. 2K, left; N = 15 cells). Taken together with the presence of ventral PIP3 patches under LatA treatment in AX4 (Taniguchi et al., 2013), the results indicate that PI3kinase and its upstream Ras are central to the patch dynamics (Fukushima et al., 2019) and they are strongly dependent on micro-topography.

The high directedness of topographically guided cell movements raises a question about their potential crosstalks with chemotaxis and the role of PI3K. Upon binding of cAMP to a G-protein coupled receptor, transient activation of PI3K takes place over the course of a few minutes. However, because null mutant of PI3Ks are still able to chemotax (Hoeller and Kay, 2007), the role of PI3K is not entirely clear. When cells undergoing topography guidance were exposed to a concentration gradient of cAMP formed from the tip of a glass needle, two distinct types of response were observed (Fig. 3A-D, Movie. S4). In approximately half of the case, F-actin patches disappeared within 5 minutes after cAMP stimulation (Fig. 3E; 37.5 % on the microstructured surfaces (N = 40 cells) and 58.5 % on the flat (N = 41 cells)), and the cells migrated up the cAMP gradient (Fig. 3A). For the rest of the case, actin patches persisted and cells continued to migrate along the ridge irrespective of the orientation of the gradient (Fig. 3B). Trajectories of representative 10 cells are shown in Fig. 3C (on flat surfaces) and 3D (on microridges), clearly indicating the distinct chemotactic behaviors according to the presence of F-actin patches. The percentage of cells with patches declined more rapidly in cells on the flat surface compared to those on the structured surface (Fig. 3E). After 10 min, patches are restored in cells on the structured surface while they remain diminished in cells on flat surfaces which further supports the inductive role of topography in the patch formation. The mean migration speed toward the cAMP source was close to zero in patch-positive cells (Fig. 3F). The same cell was observed to switch between the two behaviors; cells that first migrated up a cAMP gradient (Fig. S5A, 0-100 sec) stopped as a ventral actin patch appeared (Fig. S5B, 110-210 sec), then resumed chemotaxis as soon as the patch disappeared (Fig. S5B, 220-300 sec). In addition, small protrusions toward the cAMP source were observed in some cells during micro-topographic guidance (Fig. S5C), suggesting that patch-positive cells can still respond to extracellular cAMP, however some cells cannot override the topography guidance. While the variable responses are not unexpected considering that there are cell-cell heterogeneity in the expression level of the GPCR-signaling pathway, the erasure of the patch upon cAMP stimulation in majority of the cells and its later recovery (Fig. 3E) suggest that the transient activation of PI3K by cAMP can serve to release cells from directionality imposed by the topography at least temporarily.

**Figure 3.**
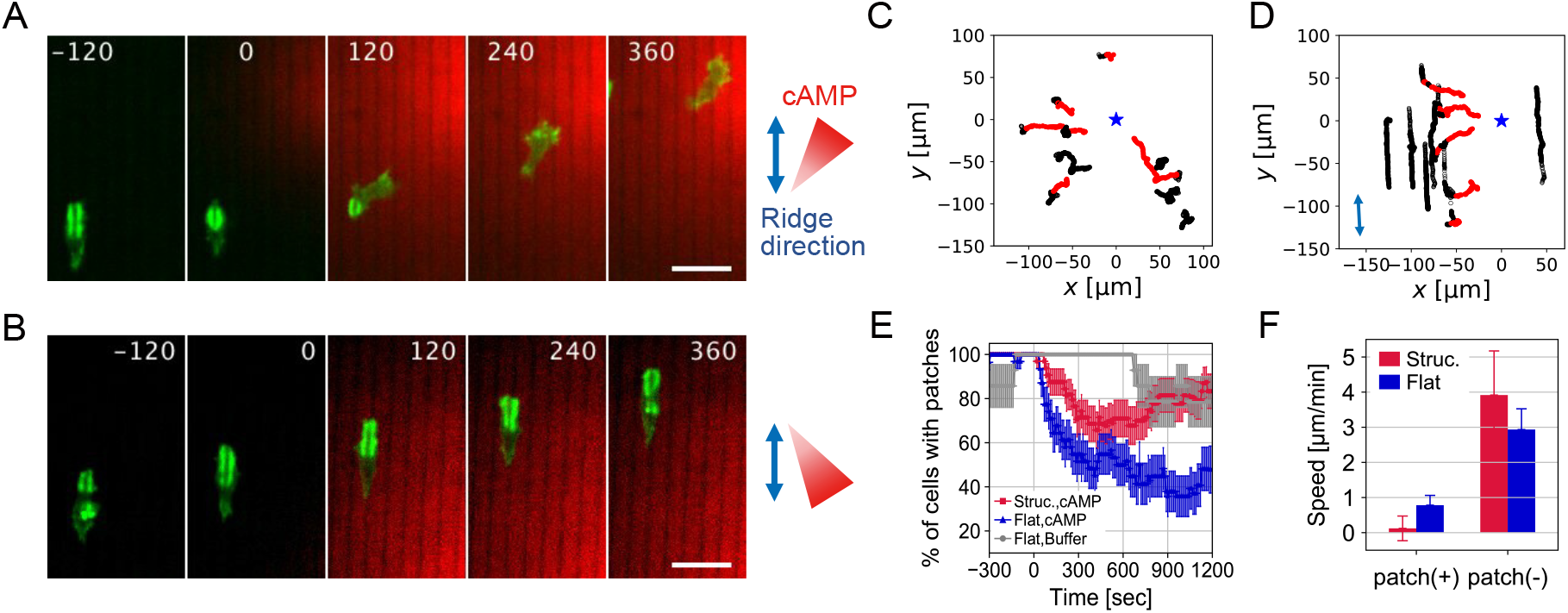
Micro-topographical guidance is independent from chemotaxis. (**A** and **B**) Confocal timelapse images of cells on the microridges (green: GFP-Lifeact fluorescence, red: Alexa594). Time in sec; cAMP application from a microneedle at t = 0. Scale bars, 20 μm. (**C** and **D**) Representative trajectories of cells on non-structured (**C**) and structured (**D**) SU-8 surfaces (open: t = −5 to 0 min, closed: t = 0 to 15 min, black: patch-positive, red: patch-negative, blue stars: the position of the cAMP source). (**E**) Fractional change in patch-positive cells after cAMP application (mean ± s.e., N = 30 (structured, cAMP), 30 (non-structured, cAMP) and 14 cells (non-structured, buffer)). (**F**) Cell migration speeds in the direction of the cAMP source (mean ± s.e., N = 36 (structured, patch(+)), 17 (structured, patch(-)), 32 (non-structured, patch(+)) and 25 cells (non-structured, patch(-))). Cell migration speed toward cAMP source was determined by quantitating the net centroid displacement towards the microneedle tip for Δ*t* = 10 sec.

### Substrate curvature determines the F-actin intensity and the propagation direction of the signaling patches

To further address geometrical features of the substrates that constrain the direction of patch propagation, we employed an SU-8 surface with a large plateau and analyzed the patch dynamics along its well-separated convex and concave corners. If the height of plateau (*h*) was large enough to prevent patches from covering both the top and bottom plane at the same time (*h* = 8.5 μm), we found that patches propagated along the convex edge (Fig. 4A). For lower plateaus (*h* = 3.5 μm), patches that traveled down from the top and reached the bottom (Fig. 4B, 00:00 ~ 01:00) did not move across the concave edge. Rather than continuing to spread across the lateral plane, it always turned in the orthogonal direction and travelled along the edge (Fig. 4B, 01:00 ~ 01:30). These observations suggest two opposing effects by the surface topography; convex surfaces attract and guide the patch, while concave surfaces prevent it from propagating further. For more rigorous quantification of this effect, reconstructed 3D confocal images were analyzed for spatial occupancies of patches with regard to the surface topography (Fig. 4C). Here, *D_U_* and *D_H_* are the distances from the convex edge to the farthest points covered by the actin patches at the top and lateral planes, respectively. *D_L_* is the distance from the concave edge to the farthest point within the patches at the bottom plane, and *D* is the sum of *D_U_*, *D_H_* and *D_L_*. We found that, for *h* = 8.5 μm, *D_U_*/*D* = 48 ± 1 % and *D_H_*/*D* = 51 ± 1 % (Fig. 4D, N = 83 plots), meaning that patches expanded equally well towards the top and lateral planes. At the intermediate height *h* = 3.5 μm, *D_U_*/*D* increased to 59.4 ± 0.9 % while *D_H_*/*D* decreased to 35.8 ± 0.7 % and *D_L_*/*D* = 4.9 ± 0.5 % (Fig. 4D, N = 117 plots). Because *D_U_*/*D* > (*D_H_ + D_L_*) / *D*, the concave edge must be inhibitory. Note that, by definition, *D_U_*/*D* = (*D*_H_ + *D*_L_) /*D* at *h* > 0 means no confinement effect. For a low plateau *h* = 1 μm, *D_L_*/*D* increased to 23.8 ± 0.6 % (Fig. 4D, N = 407 plots), reflecting the spread of the patches at the bottom plane.

**Figure 4.**
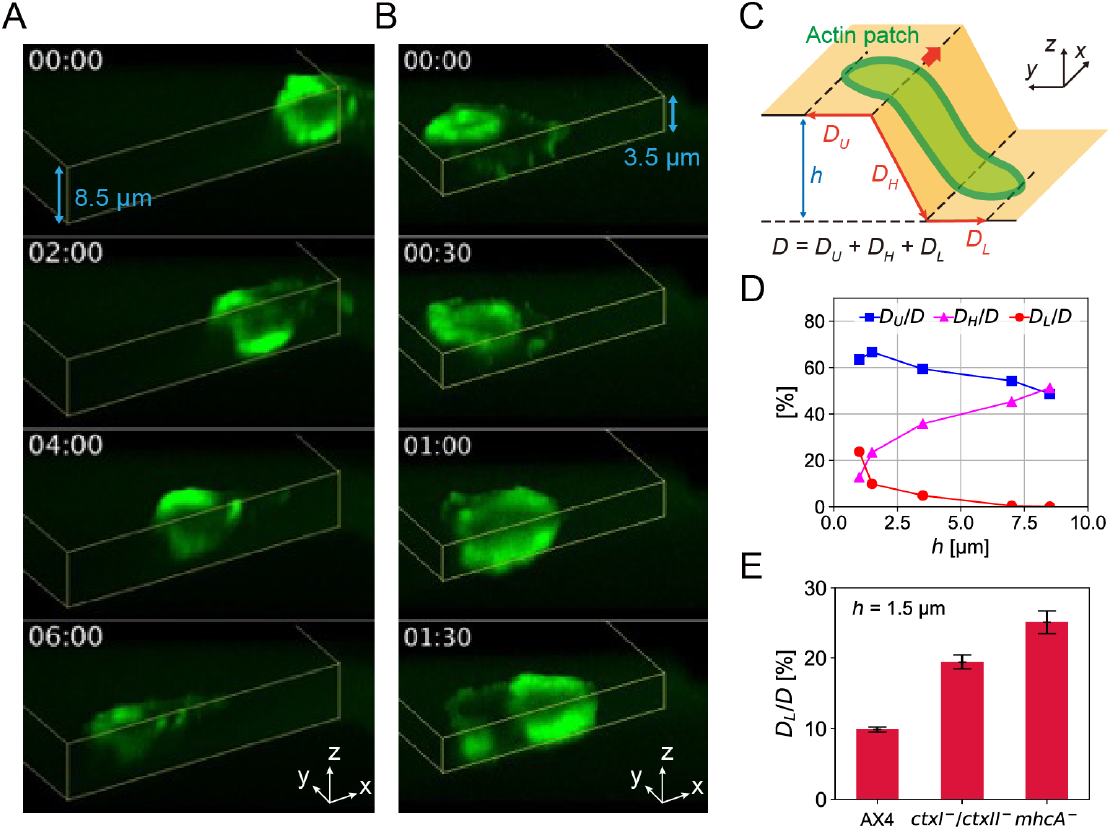
The direction of patch propagation is determined by convex and concave surfaces. (**A** and **B**) 3-D timelapse confocal images of patch-positive cells on 8.5 μm (**A**) and 3.5 μm (**B**) height plateaus (green: GFP-Lifeact fluorescence; MIP in the direction 60 degrees from the z-axis, yellow lines: contours of plateau surfaces). Time in min:sec. (**C**) A schematic for parameters *D_U_*, *D_H_*, *D_L_*, *D* and *h*. (**D**) The height-dependence of *D_U_*/*D*, *D_H_*/*D*, *D_L_*/*D* in AX4 cells. (**E**) *D_L_*/*D* at *h* = 1.5 μm in AX4, *ctxI*-/*ctxII*- and *mhcA*- (mean ± s.e., N = 5, 9, 3 cells).

The analysis above indicates that the convex edge traps the patch while the concave edge blocks it from propagating further. The near vertical contact angle between the dorsal side of membrane and the bottom plane (Fig. 1D, bottom; Fig. S3B) indicates high membrane tension. We postulated that strengthening of crosslinked actomyosin meshwork at the dorsal plasma membrane may act to suppress expansion of the F-actin patch filled with the branched actin meshwork. Since crosslinkers of cortical actin - myosin II, cortexillin I and cortexillin II are the main source of cortical tension in *Dictyostelium* (Kee et al., 2012; Reichl et al., 2008), the patches may not be prevented from traveling across the concave edge in their null mutants. In support of this notion, *D_L_*/*D* for *h* = 1.5 μm increased from 9.9 ± 0.4 % in AX4 to 19 ± 1 % in *ctxI-/ctxII-* and 25 ± 2 % in *mhcA-* (Fig. 4E). Furthermore, we found that the ventral F-actin patches in *ctxI*-/*ctxII*- and *mhcA*- were less confined to the microridge (*ctxI-/ctxII-* in Fig. 6A, 0-150, 750 sec; *mhcA-* in Fig. S6A). Notably, in *mhcA*- patches often traversed the ridge and the bottom plane (Fig. S6B, C). These observations suggest the cortical actomyosin at the dorsal side is essential for the confinement of F-actin patches to the concave edge.

Since fabrication of ridges of various curvatures in z-direction requires fine 3-D photolithography and thus technically demanding, we employed microridges of the same dimension in z-direction however with patterned ridges so as to realize various curved corners in the x-y plane. For square zig-zag patterns with alternating ±90 degrees corners (Fig. 5A, Movie. S5), we observed that the cell anterior and the underlying F-actin patch faithfully traced the zig-zag pattern. The movement can be consistently understood from the topography dependence of the F-actin patch. During turning, strong accumulation of F-actin continued along the outer corner, while it diminished at the inner corner (Fig. 5B; −90 ~ −30 sec) before it recovered as the leading edge exited the corner (Fig. 5B; 0 ~ 90 sec). Fluorescence intensities of GFP-Lifeact increased transiently at the outer corner by 1.6-fold (Fig. 5C; 90 degrees) and decreased down to 0.2-fold at the inner corner (Fig. 5D; 90 degrees). Similar experiments were then performed using ridges with corners set at ±120 degrees angle. There, the intensity fluctuations were smaller (Fig. 5C; 120 degrees), indicating that corners with sharper angles are more effective in enhancing F-actin accumulation. Because the patch can interface with more than one edge, we also tested how they respond when encountered with inconsistent corners at each side. On a ridge with T-junctions, actin patches that entered the junction from the bottom of the T faces two 90-degrees corners facing the opposite directions. There, the patch stalled at both sides of the ridge and sometimes reversed its direction (17.2 %, N = 29 events, Fig. S7A(a)). When the actin patches and the cell anterior entered the junction from the top of the T, patches continued to propagate at the straight side while they were stalled at the concave side (Fig. 5E). The percentage of patches that reversed its direction was 15.5 % (N = 84 events, Fig. S7A(b), “Reverse”). We also tested X-junctions, which also have concave corners at both sides (Fig. 5F, Movie. S6). As in T-junctions, patches stalled as cells entered the intersection (Fig. 5F; 0 ~ 180 sec), then began to propagate in the reverse direction (Fig. 5F; 180 ~ 360 sec) (27.6 %, N = 76 events, Fig. S7A(c)). In Y-junctions, reversal also occurred in 29.3% of the AX4 (Fig. S7B, “Reverse”, N = 58 events). Patch reversal at the Y-junction was never observed for *ctxI*- and *ctxI*-/*ctxII*- (Fig. S7B, “Reverse”, N = 43 and 57 events, respectively). These results further vindicate that concave surface suppresses F-actin patches and that this inhibitory effect depends on the cortical actin at the dorsal plasma membrane.

**Figure 5.**
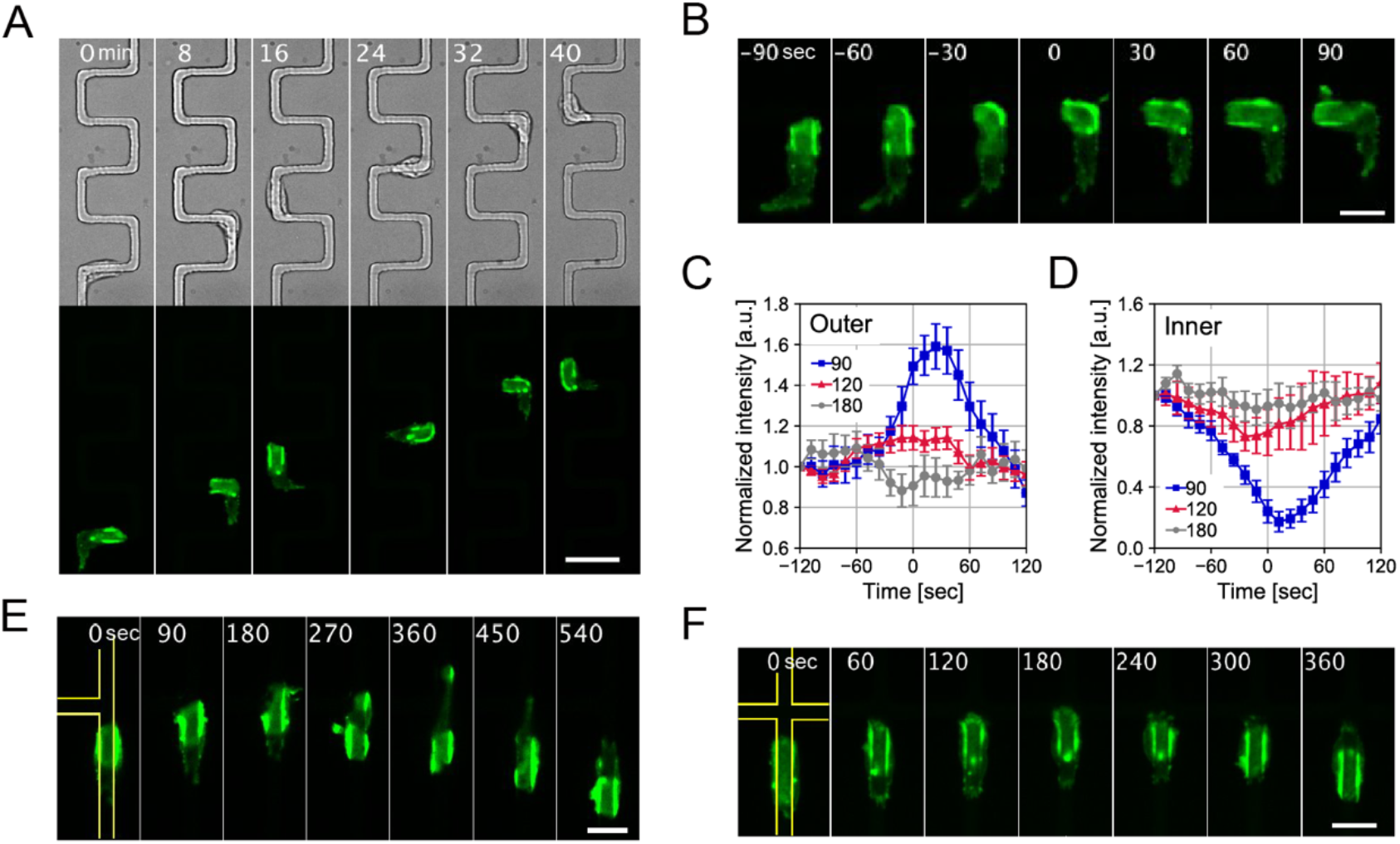
F-actin accumulation depends on the corner angle of zig-zag and bifurcating ridges. Patch and evagination guidance along microridges (*h* = 1.5 μm, width 4 μm) with corners. (**A** and **B**) Zig-zag microridges with 90 degrees corners. (**A**, upper panel) Transmitted light images. (**A**, lower panel and **B**) GFP-Lifeact fluorescence; MIP from z = 0 to 2 μm. Zoom-up images of turning along the corner (**B**). (**C** and **D**) Change in the GFP-Lifeact intensity along the outer (**C**) and inner corners (**D**). Angles are 90 (blue), 120 (red) and 180 (grey) degrees (mean ± s.e., N = 16, 21, 11 events). t = 0 is the time when patch centroid reached the corner. (**E** and **F**) Patch reversal at T-junction (**E**) and X-junction (**F**). Yellow lines indicate the ridge contours. Time in min (**A**) and sec (**B**, **E**, **F**). Scale bars, 20 μm (**A**) and 10 μm (**B**, **E**, **F**).

In addition to turning, traveling patches frequently split at the junctions (Fig. S7A, B, “Split”). For quantification, we employed the Y-junction (Fig. 6A, Movie. S7) since its three-fold symmetry made data-sampling more efficient than the T-junction. Out of all cells that entered the Y-junction, 40.0 % in AX4, 86.0 % in *ctxI-* and 75.4 % in *ctxI*-/*ctxII*- ended up splitting (Fig. S7B, “Split”). As bifurcated patches continued to propagate and extend the membrane along the respective branch (Fig. 6A, 150 ~ 450 sec), expansion of the leading edges slowed down and came to a halt when one of the competing patches disappeared (Fig. 6B, 340 ~ 600 sec). As soon as a patch diminished on one side, the cell body rapidly retracted towards the surviving branch (Fig. 6A, 600 ~ 750 sec). Despite large stretching, there was no cell division or fragmentation as observed in giant cells fused by electric pulses (Flemming et al., 2020). To gain an insight into the process of the ridge selection, the time evolution of patch size and cell elongation along the respective branch were analyzed (Fig. S7C-E). In all strains studied, AX4, *ctxI-* and *ctxI-/ctxII-,* split patches were asymmetric in size from the beginning till the end, and the larger patch survived in the majority of cases (Fig. S7D). The maximal distance from the junction point to the bifurcating cell edge (Fig. 6A, *t* = 600 inset) in the side of the surviving patch *l_s, MAX_* was always larger than those of the diminished patch *l_d, MAX_* (Fig. 6C, Fig. S7E). In *ctxI-* and *ctxI-/ctxII-*, both *l_s, MAX_* and *l_d, MAX_* were long compared to AX4 (Fig. 6C). The relative elongation (*l_s, MAX_* + *l_d, MAX_*) / *l_0_* was 1.29 ± 0.08, 1.59 ± 0.09 and 2.3 ± 0.1 for AX4, *ctxI-* and *ctxI-/ctxII-* (N = 23, 32, 36 events), respectively. *l_0_* is the front-to-back length of a cell prior to splitting. The patch was less confined to the ridge and thus spread at the bottom plane (Fig. 6A, 0 ~ 150 sec). As the patch split, they were retracted to the side of the ridge (Fig. 6A, 300 ~ 450 sec). As soon as the patch disappeared on one side, the surviving patch spread out to the bottom plane (Fig. 4A, 750 sec). The observation suggests that the elevated tension exerted by the bifurcated protrusions help confine the patch to the ridge even when cortical actomyosin is reduced. Indeed, the cell area outside the ridge in the splitting patches normalized to that in the non-splitting patches was *A_S_*/*A_NS_* = 0.68 ± 0.04 (N = 14 cells) and 0.71 ± 0.04 (N = 17 cells) in *ctxI*- and *ctxI*-/*ctxII*-, respectively compared to 1.06 ± 0.04 (N = 8 cells) in AX4 (Fig. 6D, E). These measurements suggest that the patch competes with tension-based repression which may not strictly require the presence of cross-linked actin meshworks.

**Figure 6.**
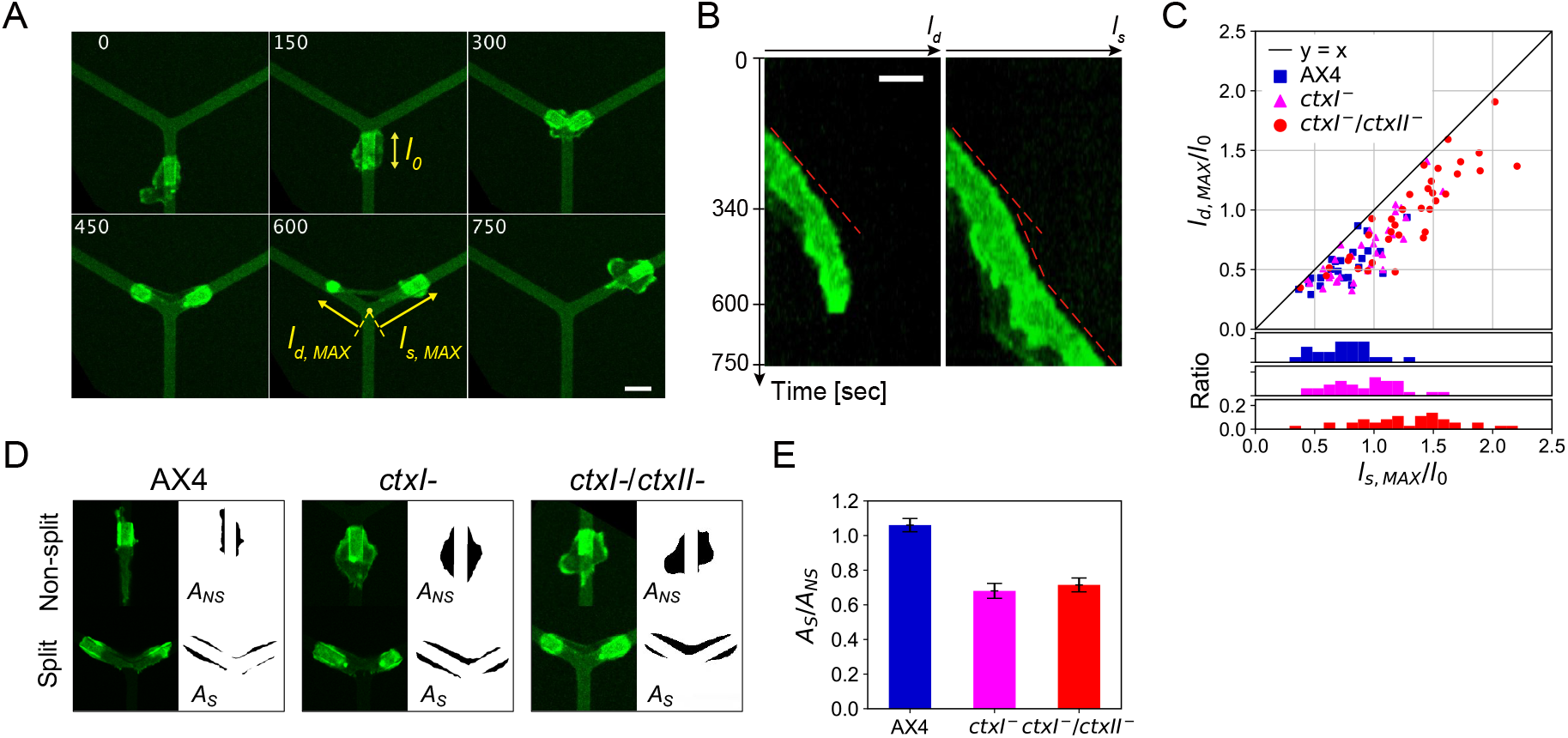
Membrane tension facilitates confinement of the patches to the ridge. (**A**) Time-lapse confocal images of Lifeact-neon/*ctxI*-/*ctxII*- at Y-junction. Time in sec. Scale bar, 10 μm. The ridge is 1.5 μm high and 4 μm wide. (**B**) Kymographs taken along each ridge branch in **A**. Images in **B** are enlarged four times in time-axis. Scale bar, 10 μm. (**C**) Scatter plots (upper panel) and histograms (lower panles) of maximal elongation rate *l_s,MAX_* /*l_0_* and *l_d, MAX_*/*l_0_* in AX4, *ctxI*- and *ctxI*-/*ctxII*- (N = 23, 32, 36 events). (**D**) Representative snapshots (left panels, green: GFP-Lifeact or Lifeact-neon fluorescence) and cell masks outside the ridge (right panels) of AX4, *ctxI*- and *ctxI*-/*ctxII*- cells with a single patch (upper) and two split patches (lower). (**E**) Ratio of *A_S_* and *A_NS_*, cell areas outside the ridge during patch splitting and otherwise, in AX4, *ctxI*- and *ctxI*-/*ctxII*- cells (mean ± s.e., N = 8, 14, 17 events).

Overall, the above results indicate propensity of the patch to position itself above convexly curved surfaces, and either be blocked or extinguished at the concavely curved surfaces. Besides the patch dynamics, we asked whether the membrane protrusion itself can capture the ridge. In immune-cells, phagocytosis involves a zippering mechanism where the advancing rim of a cup protrudes sequentially by forming an anchorage with the specific surface signal, and the membrane maintains close contact with the surface to gain traction (Jaumouillé and Waterman, 2020; Swanson and Baer, 1995). Macropinocytic cup formation, on the other hand, does not require such specific anchorage and traction (Jaumouillé and Waterman, 2020; Swanson and Baer, 1995). To see whether non-zipper type cup formation is able to capture a ridge, we tested a minimal model of macropinocytic cup formation (Saito and Sawai, 2020). The model describes a reaction-diffusion process at the plasma membrane that forms propagating signaling patches that grow in size until they consume a finite resource; e.g. total number of Ras molecules, actin nucleators, etc. The protruding force is perpendicular to the membrane, and it is restricted to the edge of a patch, which is plausible in light of the SCAR/WAVE complex localization (Bretschneider et al., 2004) and the alignment of actin filaments at this region (Jasnin et al., 2019). Although the model predicts that the geometry of the patch patterning should naturally displace the position of the SCAR/WAVE complex from the edge of the protrusion to yield the necessary inward tilt for cup formation (Saito and Sawai, 2020), other processes that could yield the inward force should serve well for the present purpose. On flat surfaces, the cups cannot form due to large load by the physical barrier (Fig. 7A-C). At the ridge, protrusions are released from the frustrated state and successfully captured the ridge (Fig. 7D-F, Movie. S8). Note that, in order to test the minimal requirement, adhesion strength between the membrane and the surface was assumed to be uniform, and no topography dependence of the patch dynamics was included. The simulations demonstrate that the primary motive force in a ring-like profile is sufficient to trap the patch and hence lock the overall cell orientation.

**Figure 7.**
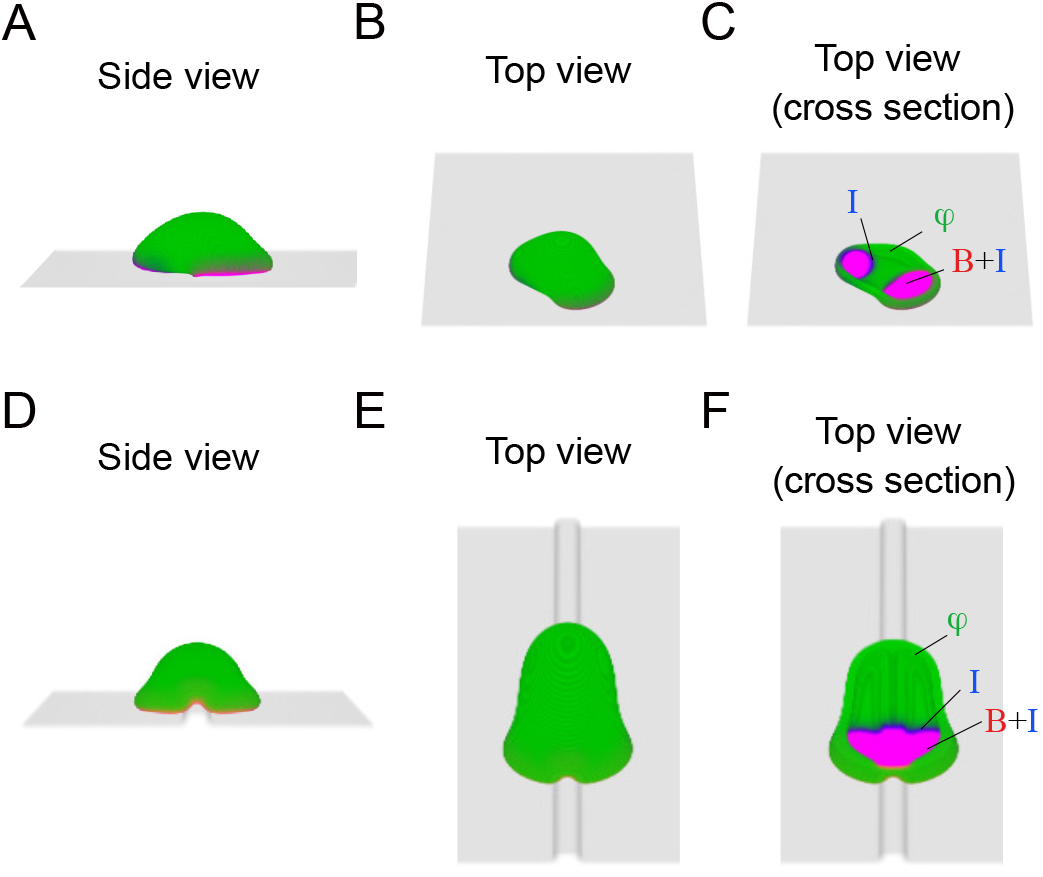
Macropinocytic cup formation can capture a microridge. (**A**-**C**) Representative snapshots of simulations on a flat substrate. The signaling active patch (red; *Aψ* > 0), the inhibitor molecule (blue; *Iψ* > 0) and the membrane (green; *ψ* > 0) shown as merged RGB images; side view (**A**), birds-eye view (**B**), the cross section along the plane parallel to the surface (**C**). (**D**-**F**) Representative snapshots of model simulations with a microridge of height *=* 1.5 μm and width = 3.0 μm. Parameters: *dx* = 0.2 μm, *dt* = 2 × 10^−4^ sec, *ϵ* = 1.2 μm, *M*_*V*_ = 5.0, *τ* = 10.0 nN ⋅ sec/μm^3^, *F* = 2.6 nN/μm^2^, *η* = 0.7 nN/μm, *β* = 100.0, *θ* = 0.105 *k*_1_ = 0.05, *k*_2_ = 0.5, *a*_*t*_ = 1.6, *D*_*a*_ = 0.17, *D*_*i*_ = 0.1 for (**A**-**C**) and *D*_*i*_ = 0.13 for (**D**-**F**), *K*_1_ = 0.05, *K*_2_ = 0.04.

## Discussion

Previous works have suggested that signaling patches of PIP3 and Ras in *Dictyostelium* are constitutive process that serve as templates for phagocytic/macropinocytic cups (Gerisch et al., 2009; Veltman et al., 2016). The present work demonstrated that the patch dynamics are topography-dependent and the resulting topography guidance serves to steer membrane protrusions along micrometer-scale convoluted surfaces. The topographical guidance was PI3K-dependent and was mediated by the following three properties: 1) F-actin independent patch initiation and growth that is selective to convex surface of micro-meter scale, 2) patch confinement at the concave surface and 3) physical capturing of the ridge by the membrane protrusion. The patch confinement was not unexpected given the fact that the ventral F-actin waves are trapped at the sidewall of perforated microwells (Jasnin et al., 2016) and that phagocytic cups are known to stall at the furrow of a budding yeast where IBARa is localized (Clarke et al., 2010). Phosphatidylinositol 3,4,5-trisphosphate 3-phosphatase PTEN is known to accumulate at the aspirated region, where the positive feedback from actomyosin potentiates its force-induced translocation (Pramanik et al., 2009). Heightened activity of PTEN may recruit more cortexillin to the plasma membrane and the resulting crosslinked actin meshwork should prevent expansion of the F-actin patch. The resulting membrane tension should form a positive feedback loop to further suppress patch propagation at the concave edge. Such notion is in line with our observation that confinement at the concave surface was reduced in the null mutants of myosin II and of cortexillins (Fig. S6, Fig.6A). The fact that the reduction was rescued when the cells were stretched at the bifurcating ridges (Fig. 6D-E) further strengthens the notion that the patch confinement is mediated by membrane tension rather than directly through cortical actomyosin. These features may be related to extinction of actin waves in neutrophils when they collide with physical obstacles (Weiner et al., 2007) and the tension-mediated suppression of the leading edge in migrating cells.

On the other hand, the topography-dependent patch nucleation and the ridge capturing suggest a novel mechanism whereby a cup is selectively generated at the plasma membrane in contact with sufficiently curved surfaces and the ability of the resulting ring-like protrusion to capture the ridge. Since a misplaced patch would have high chance of being confined to the concave surface, it is essential that a patch is selectively induced at the convex surface for efficient surface capturing. Topography guidance was completely eliminated by LY treatment (Fig. 2B-D) indicating that the patch nucleation at the ridge or the physical capturing itself or both are PI3K dependent. Since the latter requires sustained presence of PIP3 patches, contributions from the two processes is difficult to separate. *Dictyostelium discoideum* has five class-I PI3Ks, in which PI3K1/2 are essential to nucleate PIP3 patches in the early stage of macropinocytic cup formation (Hoeller et al., 2013). Activity of PI3Ks requires interaction with RasG, RasS and Rap at its Ras-binding domain (Hoeller et al., 2013; Kortholt et al., 2010). Patch of pan-RAS-GTP probe Raf1RBD has been observed in PI3K-null (Veltman et al., 2016) under LatA treatment (Fukushima et al., 2019) indicating that Ras is the central driver of the patch dynamics. Since forced elevation of Ras/Rap activity can elevate spontaneous patch generation (Miao et al., 2017), the present analysis indicates that surface topography has a similar effect on Ras/PI3K but acting locally. In the present study, topographical dependence was strictly observed in the wild-type NC4 strain (Fig. 2E-F) which is poor at liquid uptake. Even in the derivative axenic strain which has hyper Ras activity and can constitutively generate patches on flat surfaces (Bloomfield et al., 2015; Veltman et al., 2016), our results demonstrated that, when presented with microridges, patch initiation occurred exclusively at the ridges (Fig. 2E) suggesting that, topographical-sensing can operate on top of the elevated Ras/PI3K activity. Deletion of all five PI3Ks in *Dictyostelium* significantly decreases the ingestion rate for yeast particles of 3-5 μm in diameter (Chen et al., 2012) but not for bacteria or 1 μm latex beads (Hoeller et al., 2013). Interestingly, micrometer-scale dependencies are also seen in other systems. Phagocytic uptake of IgG-coated beads in RAW264.7 cells requires PI3K when beads are larger than 3 μm in diameter (Cox et al., 1999). In the frog egg extracts, the rate of actin polymerization on a PI(4,5)P2 coated glass beads is 2-3 times higher for 1 μm radius beads compared to 150-400 nm radius beads (Gallop et al., 2013). Taken together with our present findings, the coordinated engagement of Ras and PI3K in amplifying cytoskeletal signaling in a topography-dependent manner maybe a wide-spread mechanism for large particle uptake.

It should be noted that the PIP3/F-actin patches described here takes place in the order of minutes and a few micrometers in size. These features contrast with the much faster membrane curvature-dependent waves in mast cells which takes place in the order of seconds in association with F-BAR dependent nanoscale endocytosis that are synchronized in time and space (Wu et al., 2018; Yang et al., 2017). Strengthening the contact between the substrate and the plasma membrane almost completely abolished the F-actin patches both on the flat and structured surfaces (Fig. S4B). If the patch induction is based on spontaneous curvature of lipid and membrane bound molecules (Gov, 2018), such suppression is expected as strong adhesion will flatten the membrane. Although this is consistent with our observation on adhesive flat surface, the same was true for adhesive microridged surface suggesting that strongly adhesive conditions may directly be inhibitory to patch initiation, propagation or both. Despite similarities in the clutching machineries with the mammalian counterparts, *Dictyostelium* lacks real integrin and is able to gain traction without specific extracellular ligands for the adhesion complex. Cell-substrate adhesion in *Dictyostelium* is mediated by non-specific van der Waals force (Kamprad et al., 2018; Loomis et al., 2012) that is assisted by integrin-beta like adhesion protein SibA that forms a complex with a kinase Phg2 (Froquet et al., 2012), another adhesion molecule SadA, paxillin, vinculin, two talin homologues TalinA or TalinB (Tsujioka et al., 2012). Rap is required for PI3K activation as well as Ras-binding domain containing Phg2 kinase (Gebbie et al., 2004; Kortholt et al., 2006). Both Phg2 and SadA are known to be essential for phagocytosis (Fey et al., 2002; Gebbie et al., 2004). In order to mediate adhesion, SadA requires its cytoplasmic tail region that interacts with actin cross-linker Cortexillin I (Kowal and Chisholm, 2011) which is absent from the patch (Fig. S4B)(Schroth-Diez et al., 2009). One possibility is that cell-substrate adhesion at the patch is weak and that fluctuating membrane undulation is required for the feedback amplification of the SCAR/WAVE complex (Huang et al., 2013). In several mammalian cell lines, ventral F-actin waves require a cycle of integrin engagement and disengagement to ECM (Case and Waterman, 2011). Inhibition of integrin disengagement by addition of Mn^2+^ prevents wave propagation (Case and Waterman, 2011), suggesting that cell-substrate adhesion needs to be somewhat loose and within a proper range for wave generation. A weak cell-substrate adhesion is also reported at the frustrated phagocytic cup in macrophage (Barger et al., 2019).

F-actin waves appear in the leading edge of migrating neutrophils (Weiner et al., 2007) as well as in neuronal extensions during the neurite outgrowth (Katsuno et al., 2015), however their role in *Dictyostelium* migration has long been debated. Our work demonstrated a clear example of F-actin wave directed migration in the axenic strain of *Dictyostelium*. This new mode of directed cell migration which we shall refer to as ‘phagotaxis’ likely resulted from combination of ability of the aggregation-stage *Dictyostelium* to polarize while still retaining ability to form phagocytic cup. From lack of their presence in the aggregation-stage NC4 cells, migratory roles of ventral patches in *Dictyostelium* is not clear. This puzzle parallels that for large-scale macropinocytosis in *Dictyostelium* as extracellular environment that supports it is so far unknown (Kay et al., 2019). We envisage that the natural habitat that supports efficient macropinocytosis; i.e. large patch formation in non-axenic wild type is also likely to support phagotaxis. We note that vertically confined *Dictyostelium* formed phagocytic cup sideways facing a yeast particle in contact (Fig. S8A). The cup which was more persistent in the aggregation-stage cells (Fig. S8B) supported directed migration as the cell pushed the particle forward. Such phagotactic movements may help cells transfer target particles to a better location that supports ingestion. Similar mechanisms may underlie streaming migration of macrophages in contact with target cancer cells (Sharma et al., 2012). These contact-dependent migration are also reminiscent of contact activation of locomotion that supports streaming cell aggregation and cell-type dependent cell sorting (Fujimori et al., 2019) during the multicellular stage of *Dictyostelium* lifecycle. There, cell-cell contact signal is mediated by Ig-domain containing transmembrane molecule TgrB1 and C1 acting in trans between the front and back of the neighboring cells. Although TgrB1 and C1 are polymorphic genes that mediate kin-recognition in *Dictyostelium discoideum*, no phagocytic behavior has been observed between non-compatible Tgr allotype. Lack of clear Ras/PI3K activity at the contact site (Fujimori et al., 2019) suggests that TgrB1/C1 interaction is disengaged from topography dependent behavior. Given the presence of other cell-cell adhesion proteins and ECM in the later stage of development, phagotaxis may also have a role in morphogenesis.

## Materials and Methods

### Plasmids, cell strains

Dicty-codon optimized mNeon (Tunnacliffe et al., 2018) and Lifeact was inserted into G418- and Hygromycin-resistant plasmids pDM304 and pDM358 (Veltman et al., 2009) to obtain pDM304-Lifeact-neon and pDM358-Lifeact-neon. Plasmid pDGFP-MCS-Neo-CI (Faix et al., 2001) was a kind gift from Prof. Igor Weber. Plasmid pBIG-GFP-myo (Moores et al., 1996) was obtained from Dicty Stock Center. Laboratory wild-type strain AX4 and mutant strains *ctxI-* (NBRP, S00100), *ctxI-*/*ctxII-* (NBRP, S00404) and *mhcA-* (Dicty Stock Center, DBS0236379) were transformed following the standard electroporation protocol (Nellen et al., 1984). To generate PHcrac-GFP/NC4, NC4 cells washed free of bacteria were incubated overnight in HL5 medium before and after electroporation. Cells were then resuspended for selection in bacterial suspension in DB including 10 μg/ml G418. The following strains were constructed: GFP-mhcA/PH_CRAC_-RFP/Ax4, Lifeact-neon/PI3K1^N1-487^-RFPmars/Ax4, Lifeact-neon/*ctxI-*, Lifeact-mRFPmars/GFP-CtxI/*ctxI-*, Lifeact-neon/*ctxI-/ctxII-*, Lifeact-neon/*mhcA-* and Lifeact-GFP/NC4. Following strains are described previously (Fujimori et al., 2019; Taniguchi et al., 2013): GFP-Lifeact/Ax4, Lifeact-mRFPmars/Ax4, GFP-Lifeact/PH_CRAC_-RFP/Ax4, PTEN-GFP/PH_CRAC_-RFP/Ax4, CRIB_PakB_-mRFP1/GFP-Lifeact/Ax4, Lifeact-GFP/RFP-RBD_Raf1_/Ax4, HSPC300-GFP/Lifeact-mRFPmars/Ax4, PI3K1^N1-487^-RFPmars/Ax4.

### Surface fabrication

Microstructured SU-8 surfaces were prepared by standard photolithography. The microridged surface consisted of two layers of fabricated SU-8. A glass coverslip was washed with NaOH and treated with air plasma. The first layer of SU-8 (MicroChem) was spin-coated on the coverslip and baked at 65 °C for 1 min, then at 95 °C for 3 min. Photoresist SU-8 2 or SU-8 3005 were spin-coated at 2000 rpm to achieve thickness around 2 μm and 5 μm, respectively. The substrate was uniformly exposed with UV using an aligner (MA-20, MIKASA), baked at 65 °C for 1min, 95 °C for 3 min followed by 200 °C for 5 min. After cooling down to room temperature, the second layer of SU-8 was spin-coated at the same rotation speed as the first layer and baked at 65 °C for 1 min, then at 95 °C for 3 min. The first and second layer were fabricated using the same photoresist. Chrome masks (CBL4006Du-AZP, Clean Surface Technology) were patterned by a laser drawing device (DDB-201-TW, NEOARK). The substrate was UV-treated over a chrome mask and baked at 65 °C for 1 min, 95 °C for 1 min, etched using SU-8 developer and lastly hard-baked at 200 °C for 5 min. The fabricated SU-8 substrate was for one-time use.

For construction of microgrooved glass substrates, glass coverslips (MATSUNAMI, No.1, 0.13 ~ 0.17 mm in thickness) were uniformly coated with ~100 nm thick chrome using a sputtering device (E-200S, ANELVA), then the SU-8 with the microridges were attached on top which serves as a mask (1.5 μm in height and 3 μm in width at an interval of 3 μm). The substrates were immersed into chrome etchant to etch SU-8-uncovered chrome coatings. The glass substrates covered by chrome and SU-8 ridges were etched by gas plasma (Ar : O_2_ : C_4_F_8_ : CHF_3_ = 27 : 1 : 1 : 1) using a dry-etching system (NLD-5700Si, ULVAC). Residual contaminants were removed by ethanol, piranha solution and chrome etchant. Before use, the microgrooved glass substrates were washed by ethanol and NaOH.

Microfabricated SU-8 and glass surfaces were measured using AFM (Nanowizard3, JPK, now Bruker) or DektakXT (Bruker). AFM was installed in an inverted microscope (IX70, Olympus) on active vibration isolator (Herz). Cantilevers mounted on AFM were Tap300-G (BudgetSensors) and ACTA (AppNano). HS-500MG (BudgetSensors) with 500 nm step height was used as a height calibration standard for AFM measurements. Prepared SU-8 and glass substrates were set at the bottom of φ35 mm culture dish (MatTek) using a 9 × 9 mm^2^ frame seal (SLF0201, Biorad). The chamber was plasma-treated to improve wettability, using a plasma cleaner (PDC-32G, Harrick Plasma) immediately before plating cells.

### Cell preparation and timelapse imaging

Axenic strains of *Dictyostelium discoideum* were grown while shaken at 22 °C in HL5 with 60 μg/mL hygromycin B, 10 μg/mL G418 where appropriate. Lifeact-GFP/NC4 and PHcrac-GFP/NC4 cells were cultured in developmental buffer (DB) including *E. coli* B/r at OD_600_ = 6. For live-cell imaging, growing *Dictyostelium* cells were washed twice, resuspended in DB at 5×10^6^ cells/ml and shaken at 22 °C, 155 rpm for 1 hour. Cells were then pulsed with cAMP (final concentration 50 nM) every 6 min for 4.5 hours. Starved cells were plated at ~ 3×10^3^ cells/cm^2^ on a fabricated SU-8 or glass surface described above. For observation on glass surfaces (Fig. S3), adenylyl cyclase inhibitor SQ22536 (Wako) was added at a concentration of 150 μM to circumvent cell-cell agglutination facilitated by the low adhesiveness of the glass surface. NC4 cells were collected by suspending in DB, pelleted by centrifuged at 100 rcf for 3 min and resuspended in DB. The step was repeated three times to remove bacteria. Washed cells were plated at a density of ~1×10^5^ cells/cm^2^. For observation with yeast, AX2 cells expressing Lifeact-neon and yeast *Rhodotorula mucilaginosa* (NBRP, S90641) were loaded into polydimethylsiloxane (PDMS) chamber. The chamber was fabricated as previously described (Nakajima et al., 2016). Images were obtained using an inverted microscope (IX83 or IX81, Olympus) equipped with a laser confocal scanning unit (CSU-W1 or CSU-X1, YOKOGAWA) and a EMCCD camera. For cAMP and LY-loading, 20 μl of 100 nM cAMP and 100 μg/ml Alexa594, or 1 mM LY294,002 and 10 μg/ml Alexa594 were prepared in DB and loaded into Femtotips II (Eppendorf). The tip mounted on the micromanipulator (TransferMan 4r or TransferMan NK2, Eppendorf) was pressurized at 80-100 hPa using a microinjector (IM300, NARISHIGE or FemtoJet, Eppendorf). All live-cell imaging was performed at 22 °C.

### Data analysis

Image analysis was performed using ImageJ, Python and Microsoft Excel. To quantitate the relationship between cell migration and the ventral actin patches (Fig. 1E-G), confocal and transmitted-light images of GFP-Lifeact expressing cells that were acquired using 20x objective lens at 1 min intervals for 50 min time-windows (N = 6) were analyzed. To calculate the ratio of patch-positive cells, cells were manually assigned 1 or 0 according to presence or absence of the actin patches at each timepoint and averaged over all timepoints and cells (Fig. S1A). Cell trajectories were obtained by auto- or manual-tracking of cell centroids. The average speed and distribution of migratory direction relative to the ridge were calculated from the centroid displacement in moving time-window of 1 min (Fig. 1E, F). The relationship between the mean square displacement and time was fitted by the two parameters in the persistent random walk model (Dunn, 1983) to obtain the persistence time (Fig. 1G).

For quantification of spatial distribution of ventral PIP3 patches in Latrunculin A-treated cells (Fig. 2K), 3-D confocal images of PH_CRAC_-GFP expressing cells were acquired every 10 sec for > 5 minutes from *z* = 0 (the basal surface) to *z* = 3 μm at *z*-interval of 0.5 μm. From the maximum intensity projection of Z-stacks, a region occupied by PH_CRAC_-GFP patches and the cell outlines were extracted and aligned against the average cell centroid positions. The aligned binary image stacks of PH_CRAC_-GFP patches and cell outlines were cropped to 180 × 90 pixels and averaged over 892 frames from 15 cells (Struc.) and 826 frames from 14 cells (Flat). The average intensity of PH_CRAC_ patch was normalized so that the total value of all pixels is 1. Averaged cell outlines were calculated in the polar coordinate system.

To quantitate the height dependence of patch dynamics on the plateau (Fig. 4D), Z-stacked timelapse confocal images of cells expressing GFP-Lifeact were acquired every 10 sec. Images were projected into the x-axis in Fig. 4C as maximum fluorescent intensity were displayed. In the projected images, *D_U_*, *D_H_*, lengths from convex corner to the patch edges within top and lateral surfaces, *D_L_*, a length from concave corner to the patch edge within bottom surface and *D*, the sum of these three lengths, were measured at each timepoint. Ratio *D_U_*/*D*, *D_H_*/*D* and *D_L_*/*D* were averaged over all timepoints from all cells and plotted. For quantification of the relationship between the angle of the ridge corners and F-actin accumulation (Fig. 5C, D), Z-stacked time-lapse confocal images of GFP-Lifeact expressing cells plated on the zig-zag ridges were acquired every 6 or 12 sec. From kymographs of GFP-Lifeact along the inner and outer corners, fluorescent intensities within actin patch regions were extracted and integrated at each timepoint. Time frame was aligned so that patch centroids reached the corner (Fig. 5B) at time 0. The integrated fluorescence intensities were normalized to the value at t = −120 sec and averaged over all events. The membrane extension accompanied by two split patches and the patch diameters on the Y-shaped ridges were quantitated using kymographs along both branches (Fig. 6C, S7D, E). Lengths from junction-point to the two leading edges were defined as *l_s_* and *l_d_* (*s*: survived and *d*: disappeared). Cell area outside the ridge (Fig. 6D, E) was measured using binary cell-mask images created from GFP-Lifeact or Lifeact-neon fluorescence images from which the region of ridges were subtracted. The remaining area was time-averaged according to whether cells exhibited single patch (*A_NS_*) or two split patches (*A_S_*).

### Phase-field model

An abstract field variable *ϕ*(***r***) describes the cell interior region (*ϕ* = 1) and the exterior region (*ϕ* = 0) in a 3-D coordinate ***r***. *ϕ* is continuous and varies sharply at the interface with finite width characterized by the small parameter *ϵ*. To describe the interfacial dynamics, we employed the following phase-field equation

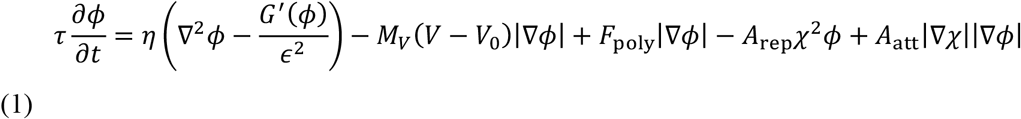

where *G*′ = 16*ϕ*(1 − *ϕ*)(1 − 2*ϕ*) and *V* = ∫ *ϕ* d***r***. The first term in the right hand side represents curvature-driven force associated with surface tension *η*. The second term describes the effective elasticity where *V*_0_ is the cell volume at the resting state and *M*_*V*_ is a fixed positive parameter. The third term describes the force normal to the interface driven by actin polymerization. The magnitude of force *F*_poly_ is a function of the local concentrations of signaling molecules as described below. The interactions between the cell and the substrate are described in the fourth and fifth term. The fourth term is the volume exclusion, and the fifth term describes the effective adhesion. The microridged substrate is described by another field variable *χ*(***r***) as follows:

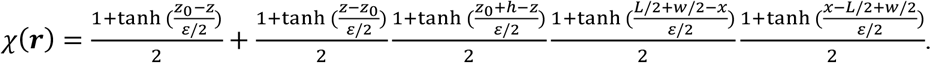

Parameters are: the offset of the substrate (*z*_0_), width (*w=*3.0 *μ*m) and height (*h=*1.5 *μ*m) of the ridge and the length in the x-direction of simulated space *L =* 40 *μ*m. The length of the simulated space in the y-direction was 60 *μ*m.

For time development of the signaling molecule, we adopted the following reaction-diffusion equations of the activator molecule (*A*) with limited total resource of molecule (*A*_t_):

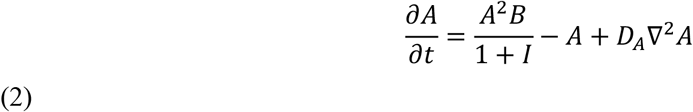

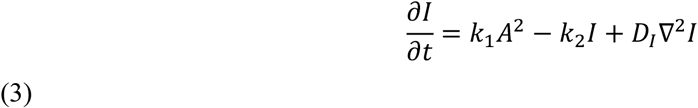

where *D*_*A*_ and *D*_*I*_ are diffusion constants of *A* and *I* molecules, respectively. In the first equation, *B* represents inactive form of the activator molecule, which is assumed to diffuse sufficiently fast and thus can be written as *B* = *A*_t_/*S* − 〈*A*〉, where *S* is the cell surface area *S* = ∫ *ψ*/*ϵ* dr^3^ and 〈*A*〉 is the total of *A* divided by *S*.

To describe the plasma membrane region, we introduced an auxiliary phase-field *ψ* = (1 + *e*^−*β*(*ϕ*(1-*ϕ*)-*θ*)^)^−1^ which defines the interface between cell exterior (*ϕ* = 0) and interior (*ϕ* = 1) region. By definition, *ψ* = 1 represents the cell membrane and *ψ* = 0 elsewhere. A sufficiently large value of *β* was chosen so that the interface is sharp. Small offset *θ* is given to render *ψ* non-zero at the interface. Using *ψ*, we arrive at the following equations:

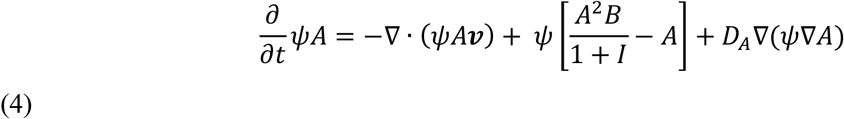

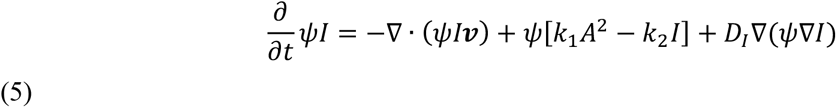

where the first terms in the right hand side are the advection term and ***v*** is given by

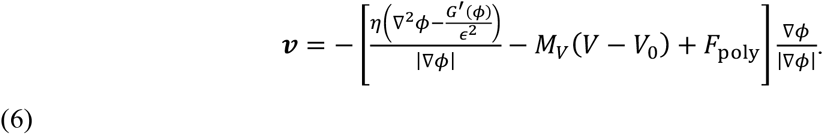

Around the substrate, Eq. (4) is given additional noise term at rate *λ* per volume. The spatial profile of the noise is given by 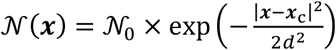, where *d* is the initial nucleation size, and 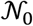 is the noise intensity that follows an exponential distribution with the average *σ*.

We assume that the magnitude of protrusion force in Eq. (1) and (6) is facilitated by *A* but attenuated by *I* following

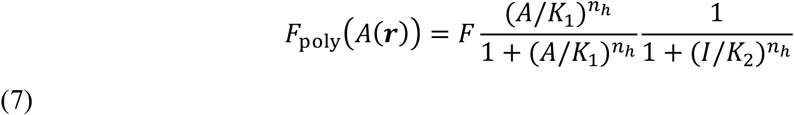

All numerical calculations were coded in C++ and run using GP-GPU GeForce GTX 1080 Ti (Saito and Sawai, 2020).

## Author contributions

GH and SS conceived the work, planned and managed all aspects of the project. GH performed all microfabrication, cell preparation, microscopy observations, and image analysis. TF, SS, and HH performed pilot experiments. GH, TF, AN, HH generated transformed cell lines. HY, TO, ST and AN assisted with microfabrication. NS formulated the mathematical model and performed numerical simulations. GH and SS interpreted data and wrote the manuscript.

## Acknowledgements

The authors thank present and past members of the Sawai lab for various technical and scientific inputs. We thank Prof. Igor Weber for pDGFP-MCS-Neo-CI, Prof. Jonathan Chubb for Dicty-codon optimized NeonGreen, Prof. James Spudich and Dicty Stock Center for pBIG-GFP-myo (DSC ID: 381), Prof. Hidekazu Kuwayama and the National BioResource Project (NBRP) Nenkin for *ctxI-* (NBRP, S00100), Prof. Günther Gerisch, Dicty Stock Center and NBRP for *ctxI-*/*ctxII-* (NBRP, S00404) and Prof. Douglas Robinson and Dicty Stock Center for *mhcA-* (DBS0236379). Takehiko Oonuki for the GFP-mhcA/PH_CRAC_-RFP/Ax4 cell line, Toyoko Sugita and Nao Shimada for pDM304-MCS-neon and pDM358-MCS-neon constructs. This work was supported by grants from Japan Science and Technology Agency (JST) CREST JPMJCR1923, Japan Society for Promotion of Science (JSPS), Ministry of Education, Culture, Sports, Science and Technology (MEXT) KAKENHI JP19H05801 to SS, Platform for Dynamic Approaches to Living System from MEXT and Japan Agency for Medical Research and Development (AMED) and in part by Joint Research by Exploratory Research Center on Life and Living Systems (ExCELLS) Grant 18-204, MEXT KAKENHI JP19H05416, JP18H04759 and JP16H01442; JSPS KAKENHI JP17H01812 and JP15KT0076 (to S.S.). GH was supported by JSPS Fellowship Grant JP18J14678.

---

**Figure S1.**
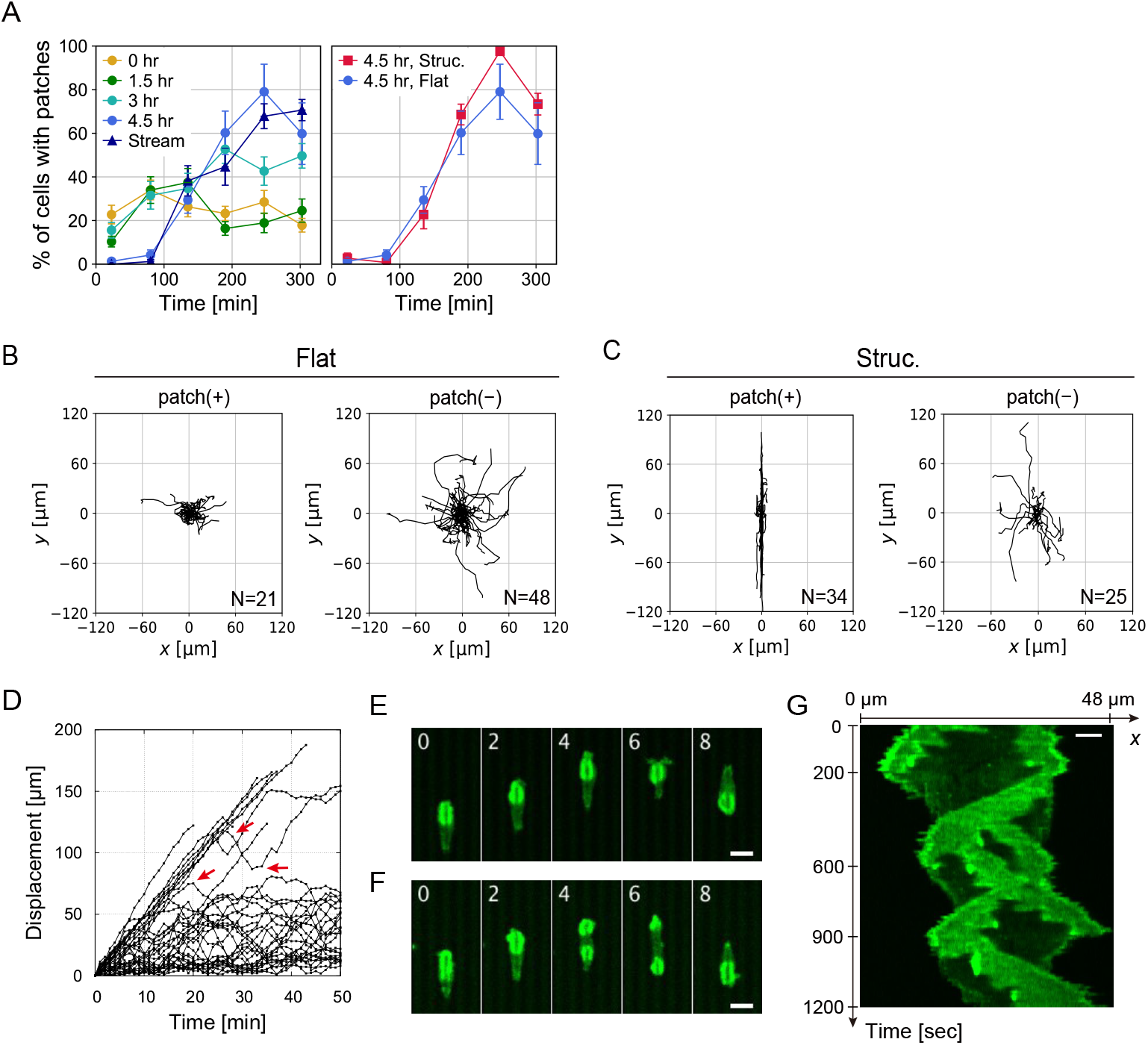
Patch nucleation and migration on flat and micro-structured SU-8 surfaces. (**A**) Change in the percentage of patch-positive AX4 cells placed on the SU-8 surface over the course of 6 hrs from the time of plating (mean ± s.e., > 17 cells per condition). (**A**, left panel) Cells on the nonstructured surface that were differentiated with cAMP pulsing (0, 1.5, 3.0, 4.5 hr) or collected from aggregation streams on agar (stream). (**A**, right panel) Cells on microstructured and non-structured surfaces that were differentiated with cAMP pulsing for 4.5 hr. (**B** and **C**) Trajectories for 20 min of patch-positive or negative cells on non-structured (**B**) and microstructured (**C**) surfaces where 1 μm high and 3 μm wide ridges were placed at an interval of 3 μm. N = 21 (Flat, patch(+)), 48 (Flat, patch(−)), 34 (Structured, patch(+)) and 25 cells (Structured, patch(−)). **(D**) Time courses of cell displacement along straight ridges in the presence of patches (N = 36 cells). Turnings (red arrows). (**E** and **F**) Confocal timelapse images (green: GFP-Lifeact fluorescence; *z* = 0). Time in min. Scale bars, 10 μm. (**G**) A kymograph taken from a cell on the ridge with small net displacement. Scale bar, 5 μm.

**Figure S2.**
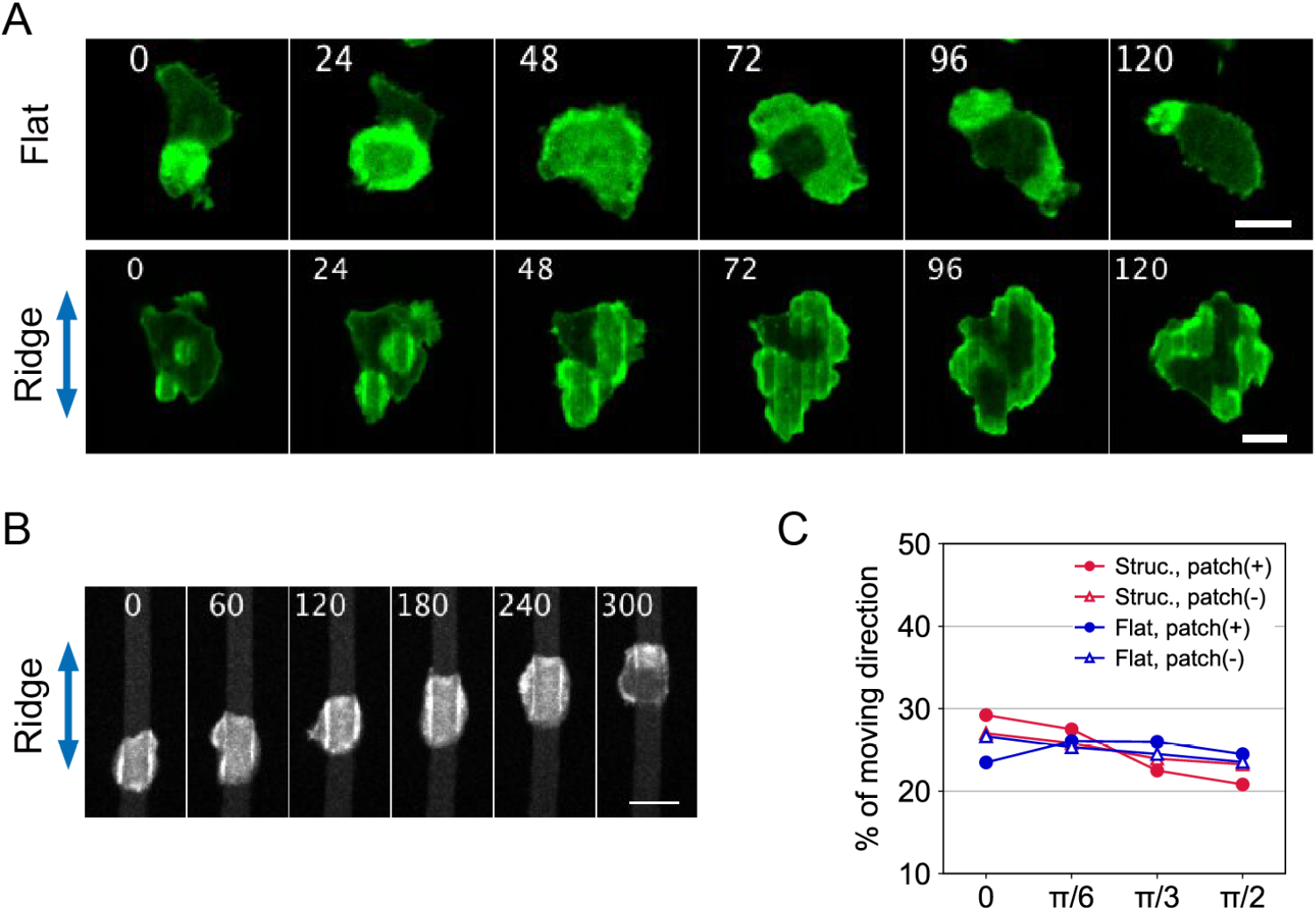
Guidance of the ventral F-actin patches and membrane evagination along a microridge in the growth-stage AX4 cells. (**A** and **B**) Time-lapse confocal images of growth-stage AX4 cells taken near the substrate. A cell on the non-structured (**A**, upper panel) and microstructured (**A**, lower panel) SU-8 surface. Ridges are 1 μm high and 3 μm wide and placed parallely at an even 3 μm spacing (green: GFP-Lifeact fluorescence; **A**, upper panel: *z* = 0, **A**, lower panel: MIP from *z* = 0 to 2 μm). (**B**) Time-lapase confocal images of a GFP-Lifeact/AX4 cell on a 1.5 μm high and 5 μm wide microridge placed parallelly at 10 μm spacing (grey: Lifeact-mRFPmars fluorecence, MIP from *z* = 0 to 2 μm). Time in sec. Scale bars, 10 μm. (**C**) Angular distribution of cell displacement relative to the ridge orientation.

**Figure S3.**
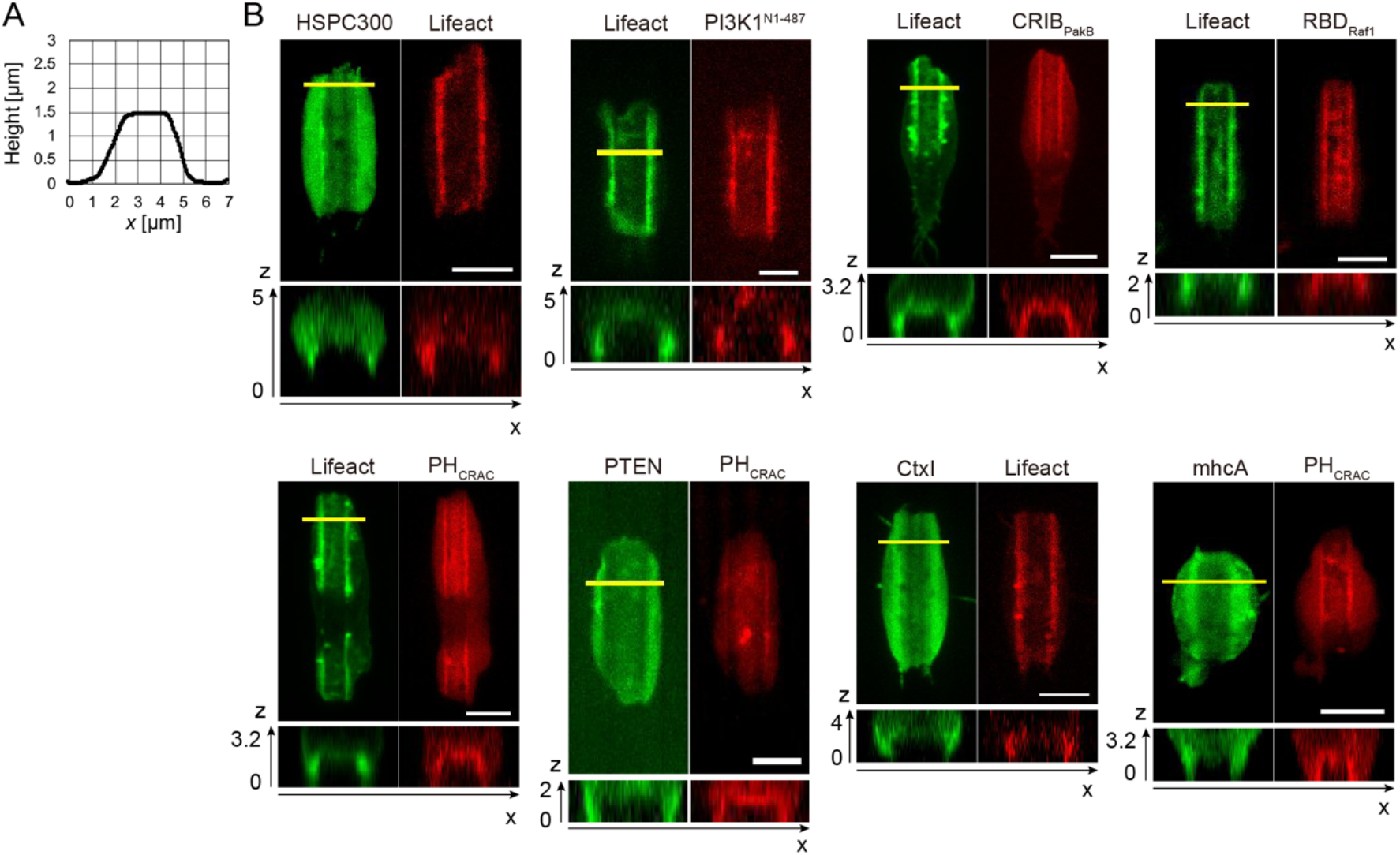
Localization of the macropinocytic/phagocytic patch components along the microridge. (**A**) The profile of an etched glass surface. (**B**) Confocal images of patch-positive cells that co-express GFP- and RFP-fluorescent probes (from left to right and top to bottom): SCAR complex/F-actin (HSPC300-GFP/Lifeact-mRFPmars/AX4), F-actin/PI3K1 (GFP-Lifeact/PI3K1Nterm-RFPmars/AX4), F-actin/Rac-GTP (GFP-Lifeact/CRIB_PakB_-mRFP1/AX4), F-actin/Ras-GTP (Lifeact-GFP/RFP-RBD_Raf1_/AX4), F-actin/PIP3 (GFP-Lifeact/PH_CRAC_-RFP/AX4), PTEN/PIP3 (PTEN-GFP/PH_CRAC_-RFP/AX4), CortexillinI/F-actin (GFP-CtxI/Lifeact-mRFPmars/*ctxI-*), MyosinII/PIP3 (GFP-MhcA/PH_CRAC_-RFP/AX4). MIP (upper panel) and an *xz*-cross section (lower panel) taken along the yellow line in the upper panel. Scale bars are 5 μm and the *z*-axis is in the unit of μm.

**Figure S4.**
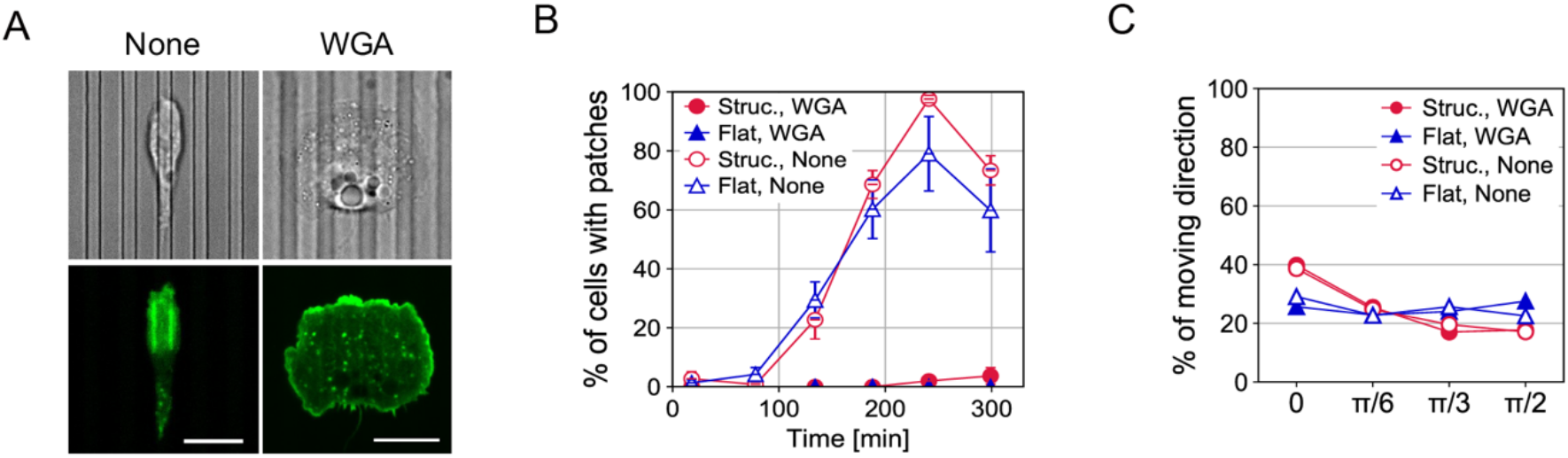
Adhesive surface suppresses ventral F-actin patches. (**A**) Aggregation-stage GFP-Lifeact/AX4 plated on the lectin WGA-coated structured substrates (upper: transmitted-light, lower: GFP-Lifeact fluorescence). Microridges are 1 μm high and 3 μm wide. (**B**) Time series of the percentage of patch-positive cells on the coated substrates (mean ± s.e., 18 cells per condition). Non-coat data are shown for comparison (duplicated from Fig. S1A, right panel). Cells are plated at time 0 min. (**C**) Angular distribution of migratory directions of cells on the coated substrates. Non-coated data are shown for comparison (duplicated from Fig. 1E, patch(-)). N = 173 (Structured, WGA), 160 (Flat, WGA), 21 (Structured, Non-coated) and 36 cells (Flat, Non-coated).

**Figure S5.**
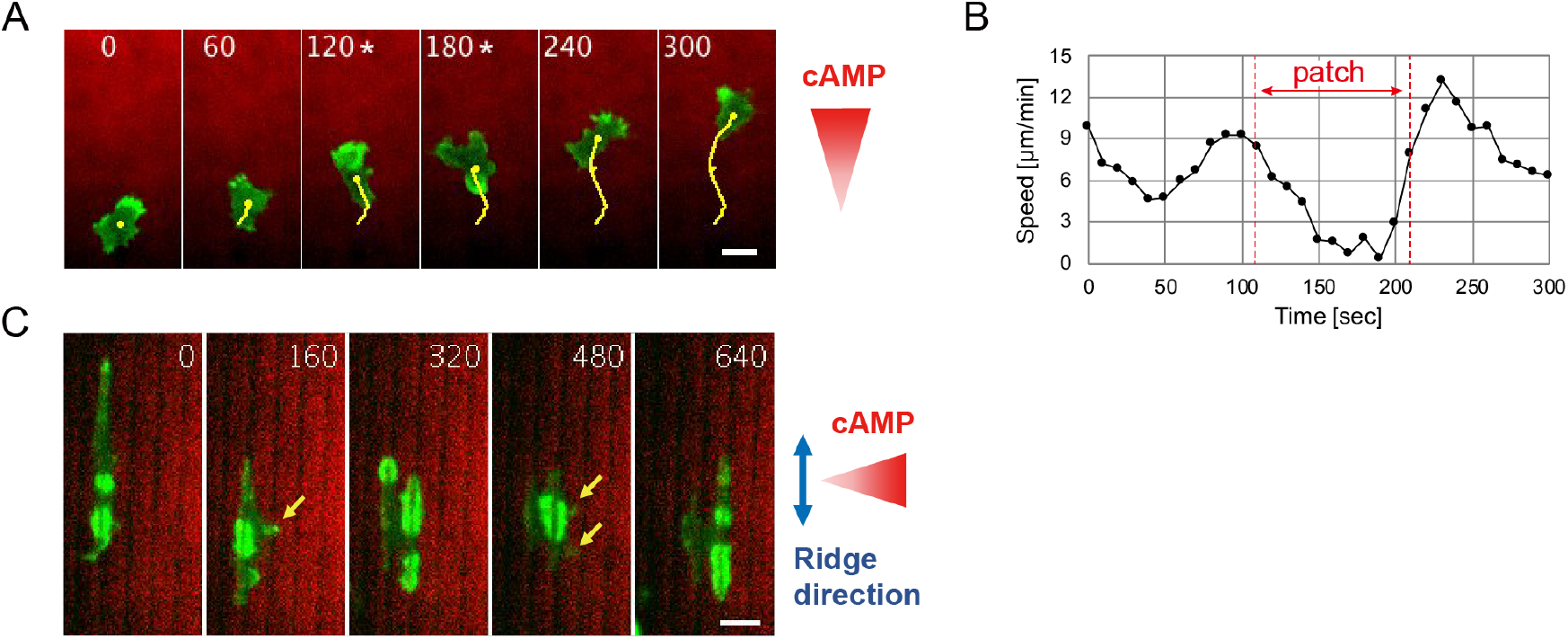
AX4 cell movement under an extracellular cAMP gradient is interfered by ventral actin patches. Aggregation-stage GFP-Lifeact/AX4 stimulated with 100 nM cAMP (green: GFP-Lifeact, red: Alexa594). Alea594 is included in the cAMP source as an indicator. (**A**) Representative time-lapse images on a flat SU-8 surface. Trajectory of the cell centroid (yellow lines). The asterisk indicates the presence of a patch. (**B**) Cell migration speed for the time sequence shown in **A**. (**C**) Representative time-lapse images of GFP-Lifeact/AX4 cells on the structured substrate. Small projections toward the cAMP source (yellow arrows). Time in sec. Scale bars, 10 μm.

**Figure S6.**
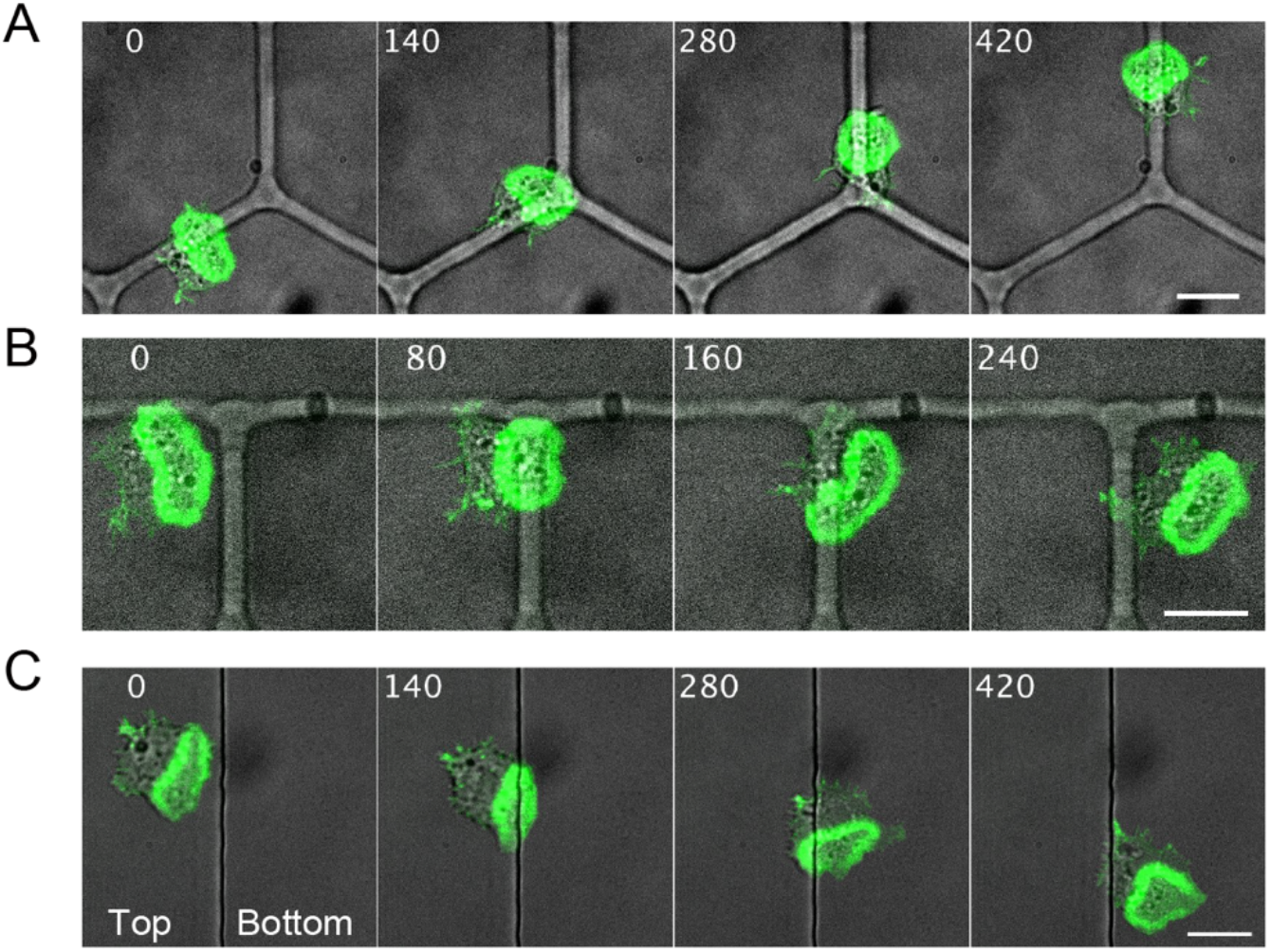
Myosin II-null cells are less confined to micro-structures. Lifeact-neon/*mhcA-* cells taken near the substrate (green: Lifeact-neon fluorescence, grey: transmitted-light). The ridge is 1.5 μm high and 4 μm wide (**A** and **B**) and the edge of a plateau surface is 1.5 μm in height (**C**). Images in **C** are MIP from z = 0 to 2 μm. Time in sec. Scale bars, 10 μm.

**Figure S7.**
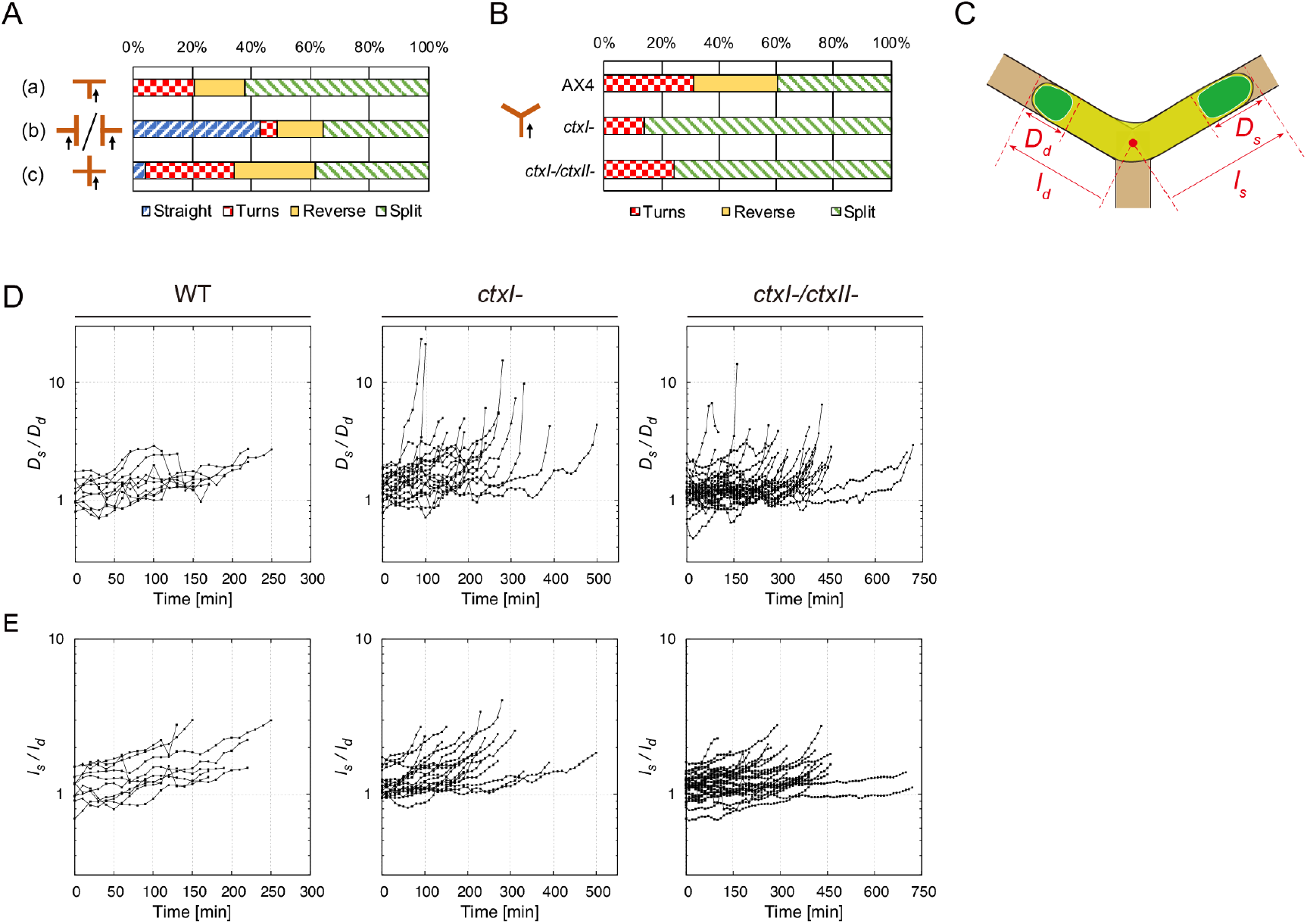
Classification and measurements of patch dynamics at the junctions. (**A**) Distribution of the patch behaviors at T- and X-junctions (a-c). N = 29 (a), 84 (b) and 76 (c) events. (**B**) Distribution of the patch behaviors at Y-junction. N = 58 (AX4), 43 (*ctxI-*) and 57 (*ctxI-/ctxII-*) events. **(C**) A schematic of patch diameters *D_s_* and *D_d_* and the distances from the junction point to the bifurcating cell edge *I_s_*, *I_d_*. Subscripts ‘*s*’ and ‘*d*’ signify patches that ‘survived’ or ‘disappeared’, respectively. (**D** and **E**) Time course of *D_s_*/*D_d_* and *l_s_*/*l_d_* at Y-junction in AX4 (N = 10 events), *ctxI-* (N = 20 events) and *ctxI-*/*ctxII-* (N = 28 events).

**Figure S8.**
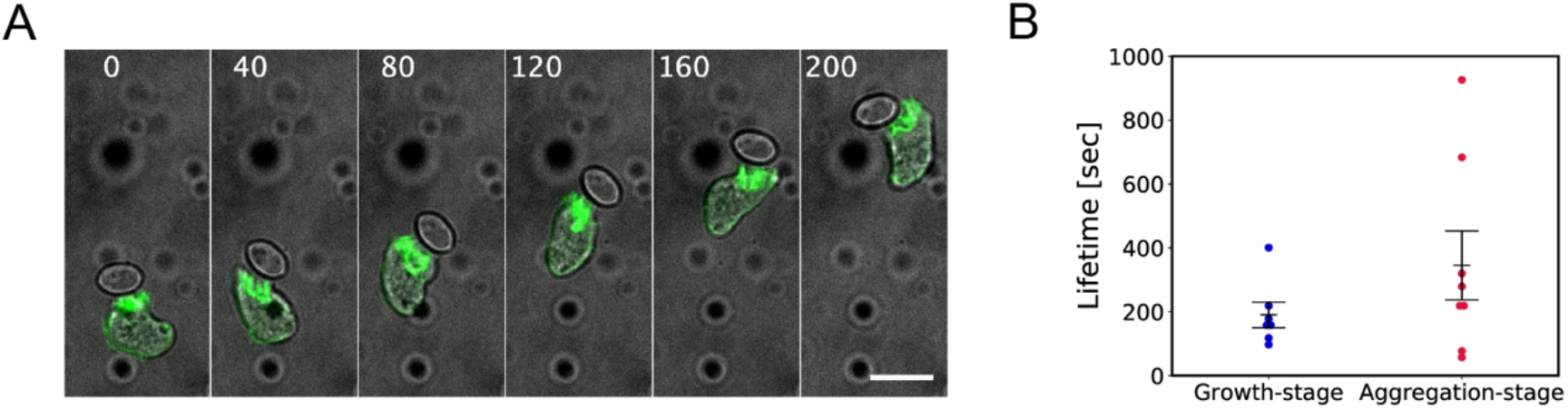
Directed migration towards an attached yeast particle. Time-lapse images of aggregation-stage Lifeact-neon/AX2 cell in contact with a yeast particle (grey: transmitted light, green: Lifeact-neon fluorescence). Time in sec. Scale bar, 10 μm. (**B**) Lifetime of actin patches at the interface between the yeast particle and *Dictyostelium* in growth-stage and aggregation-stage cells (mean ± s.e.; N = 7, 8 patches, each dot represents a unique cell).

## Movie legends

**Movie S1. Propagating ventral F-actin patch in aggregation-stage *Dictyostelium* AX4.**

Time-lapse confocal images of GFP-Lifeact/AX4 cells on a non-structured SU-8 surface. Images were acquired every 2 sec. Time in sec. Scale bar is 10 μm.

**Movie S2. Topography-guidance of the ventral F-actin patch and membrane evagination in aggregation-stage *Dictyostelium* AX4.**

Time-lapse confocal images of GFP-Lifeact/AX4 on a microstructured SU-8 surface (green: GFP-Lifeact; maximum intensity projection from z = 0 to 2 μm taken at an interval of 0.5 μm from the SU-8 surface). Images were acquired every 10 sec. The ridge is 1 μm high and 3 μm wide placed at an interval of 3 μm. Time in sec. Scale bar is 10 μm.

**Movie S3. Topography-dependence of PIP3 signaling patch in LatA-treated NC4 cells.**

Time-lapse confocal images of vegetative PH_CRAC_-GFP/NC4 cells treated with 3 μM Latrunculin A on microstructured (left) and non-structured (right) surfaces (green: PH_CRAC_-GFP fluorescence; MIP from *z* = 0 to 3 μm every 0.5 μm, the lower schematic indicates ridge positions). Images were acquired every 10 sec. The ridge is 1 μm high and 3 μm wide placed in parallel at an interval of 3 μm. Time in sec. Scale bars are 5 μm.

**Movie S4. Extinction of F-actin patch and chemotaxis to extracellular cAMP.**

Time-lapase confocal images of GFP-Lifeact/AX4 on a microstructured SU-8 surface (green: GFP-Lifeact; z-slice near the SU-8 surface, red: Alexa594, an indicator of cAMP). Transient F-actin patches (left) and a persistent patch (right). Images were acquired every 10 sec. cAMP was applied at *t* = 0 sec from a microneedle. Time in sec. Scale bars 20 μm.

**Movie S5. Guidance of F-actin patch and membrane evagination along the zig-zag patterns.**

Timelapse images of GFP-Lifeact/AX4 cells on a microridge with alternating ±90 degrees corners. Transmitted light (left) and confocal (right). Grey: transmitted light. Green: GFP-Lifeact; maximum intensity projection from z = 0 to 2 μm taken at an interval of 0.5 μm from the SU-8 surface. The ridge is 1.5 μm high and 4 μm wide. Images were acquired every 6 sec. Time in min. Scale bar 10 μm.

**Movie S6. Turning of the ventral F-actin patch and the leading edge at X-junctions.**

Confocal timelapse images of GFP-Lifeact on the microstructured SU-8 surface (green: maximum intensity projection from z = 0 to 2 μm taken at an interval of 0.5 μm from the SU-8 surface). Images were acquired every 12 sec. The ridge is 1.5 μm high and 4 μm wide. Two X-junctions appear at the top and bottom of the frame. Time in sec. Scale bar 5 μm.

**Movie S7. Splitting of the ventral F-actin patch and the leading edge at Y-junction.**

Confocal timelapse images of a Lifeact-neon/*ctxI-*/*ctxII-* cells on a microstructured SU-8 surface (green: z-slice near the SU-8 surface). Images were acquired every 10 sec. The ridges are 1.5 μm high and 4 μm wide and connected to form Y-junctions. Time in sec. Scale bar is 10 μm.

**Movie S8. Model simulations of the patch dynamics and the resulting plasma membrane deformation.**

Timelapse of the representative data shown in Fig. 7D-F for a flat surface (left panels) and for a microridge (right panel). Periodic boundary conditions are employed at *x* = 0, 40 μm and *y* = 0, 60 μm.

